# High Dimensional Proteomic Multiplex Imaging of the Central Nervous System Using the COMET™ System

**DOI:** 10.1101/2025.02.14.638299

**Authors:** Hinda Najem, Sebastian Pacheco, Jillyn Turunen, Shashwat Tripathi, Alicia Steffens, Kathleen McCortney, Jordain Walshon, James Chandler, Roger Stupp, Maciej S. Lesniak, Craig M. Horbinski, Dan Winkowski, Joanna Kowal, Jared K. Burks, Amy B. Heimberger

**Affiliations:** Department of Neurological Surgery, Northwestern University, Chicago, IL; Department of Pathology Feinberg School of Medicine, Northwestern University, Chicago, IL; Malnati Brain Tumor Institute of the Robert H. Lurie Comprehensive Cancer Center, Feinberg School of Medicine, Northwestern University, Chicago IL, 60611, USA; Visiopharm, Horsholm, Denmark; Lunaphore, Tolochenaz Switzerland; Department of Leukemia, The University of Texas MD Anderson Cancer Center, Houston, Tx, 77030

**Author notes:** Co-corresponding authors **Correspondence may be addressed to:** Amy B Heimberger, M.D, Ph.D.; Northwestern University; SQ6-516; 303 E. Superior Street Chicago, IL 60611. Jared K. Burks, PhD; The University of Texas MD Anderson Cancer Center; 1515 Holcombe Blvd, Houston, TX, 77030.

## Abstract

Sequential multiplex methodologies such as Akoya CODEX, Miltenyi MACSima, Rarecyte Orion, and others require modification of the antibodies by conjugation to an oligo or a specific fluorophore which means the use of off-the-shelf reagents is not possible. Modifications of these antibodies are typically performed via reduction chemistry and thus require verification and validation post-modification. Fixed panels are therefore developed due to various limitations including spectral overlap that creates spectral unmixing issues, steric hindrance, harsh antibody removal, and tissue degradation throughout the labeling. As such, a complex interrogation evaluating multiple study hypotheses and/or endpoints requires the development of sequential panels, reconstruction, and realignment of the tissue that necessitate a z-stack strategy. Standardized antibody panels are typically fixed and require substantial validation efforts to modify a single target and thus do not evolve with the pace of research interests. To increase the throughput of profiling cells within the human central nervous system (CNS), we developed and validated a CNS-specific library with an associated analysis platform using the newly developed Lunaphore COMET^TM^ platform. The COMET^TM^ is an automated staining/imaging instrument integrating a reagent deck for staining buffers and off-the-shelf label-free primary antibodies and fluorophore-labeled secondary antibodies, which feed into a circular plate holding up to 4 slides that are automatically imaged in microscope-operated control software. For this study, standard formalin fixed paraffin embedded histology slides are used. However, the COMET is capable of imaging fresh-frozen samples using specialized settings. Our methodologies address an unmet need in the neuroscience field while leveraging prior developmental efforts in the domain of immunology spatial profiling. Cataloging and validating a large series of antibodies on the COMET™ along with developing CNS autofluorescence management strategies while optimizing standard operating procedures have allowed for the visualization at the subcellular level. Forty analytes can be used to analyze one specimen which has clinical utility in cases in which the CNS can only be sampled by biopsy. CNS biopsies, depending on the anatomical location, can have limited available volume to a degree that requires prioritization and restriction to select analysis. In-depth bioinformatic imaging analysis can be done using standard bioinformatic tools and software such as Visiopharm®. These results establish a general framework for imaging and quantifying cell populations and networks within the CNS while providing the scientific community with standard operating procedures.

## Main

Spatial biology analysis using proteomic multiplexed imaging allows for the simultaneous analysis of several biomarkers while preserving tissue morphology [1]. Investigating the spatial architecture of a tissue broadens our understanding of cell interactions and signaling within the tissue microenvironment (TME), as well as disease pathology, progression, and treatment response. The informational yield of spatial immunofluorescence can be enhanced with other profiling modalities such as spatial transcriptomics, single-cell sequencing, and machine and deep-learning-based tools. These strategies apply to a wide variety of disciplines such as immunology, infectious disease, and neuroscience [2–4]. This type of analysis was pioneered in immune oncology because it provided assessments of the activation state and localization of immune cells within the tumor, which can be predictive of response or resistance to targeted therapeutic strategies [5–8]. As such, spatial biology analysis can impact the research pathway from initial target identification to the evaluation of clinical outcomes.

Multiplex imaging can quantify numerous proteins in tissues while retaining both morphological and spatial information. A key challenge is that a single marker does not typically define cell lineage, and multiple analytes must be labeled. The problem is further confounded when ascertaining activation states and cellular interactions. Multiplex imaging technology has been developed to overcome conventional immunofluorescence protocols, which are limited to four or five analytes per section, and a wide variety of multiplex strategies have been devised [9–25]. Imaging mass cytometry [26] and multiplex ion beam imaging [27] can assess over 40 targets using antibodies conjugated to metal reporters; however, these methods are technically challenging and expensive. As such, they have not achieved widespread scientific adoption. The Nanostring GeoMx platform [28] functions much like laser capture microdissection, yet it can detect >100 protein targets conjugated to oligonucleotide barcodes. Of note, these data are not spatial images but rather spatially registered counts of released oligos from regions of interest controlled by the researcher [1]. As such, these data are not able to generate cellular-level details. Similar limitations exist for the other transcriptomic-focused platforms. Two newer systems, Xenium and CosMX, require customized antibody conjugation limiting the research possibilities. Ultimately, most of these platforms are not able or are limited in their ability to assess biology at the single-cell level including direct cellular interactions with relevant statistical replicates. Notably, these platforms are not widely available to all scientific communities.

To address some of these limitations, we found that a microfluidic-based staining system [29–35] that performs automated staining, imaging, and elution cycles could markedly increase the ease and ability to interrogate the number of analytes analyzed in any given specimen. In a single cycle, two analytes can be detected using off-the-shelf unconjugated primary antibodies, followed by any commercially available fluorescently labeled secondary antibodies. Once the fluorescent signals are acquired, the antibody complexes are removed during an elution step, readying the section for the labeling and detection of another set of analytes. As such, sample manipulation is not required between staining steps. At the end of the full protocol, automated image registration is conducted allowing the generation and analysis of the assembled images immediately. A fully automated workflow completes the multiplexed immunofluorescent staining with a throughput of 30, 20, and 12 slides per week for 10-, 20-, and 40-plex panels per each, respectively [35]. Herein, we provide a validated inventory of antibodies with optimized protocols alongside fluorescence signal thresholding and cycle placement for the COMET™ system that can be utilized in the neuroscience field. These results should be a valuable resource utilizing a system that is commercially available and sufficiently easy to use that it will likely have widespread adoption.

## RESULTS

### Technology Platform

A pre-commercial (Priority Access or PA) Lunaphore Technology COMET™ unit was installed at Northwestern University as benchtop equipment to perform sequential immunofluorescence (seqIF™) technology for *in situ* protein characterization that involved projects spanning requirements for both discovery and validation. During the evaluation period, the COMET™ generated data for over 50 research projects spanning a variety of biological and pathophysiological conditions. Several hundred slides were analyzed and optimized for more than 200 markers in the domain of neuroscience and brain tumors (**Supplementary Table 1**). Thereafter, we transitioned to the commercial unit.

### Tissue acquisition

Under the Northwestern University Institutional Review Board approved protocol STU00214485, *en bloc* surgically resected tumors including adjacent normal human brain (**Fig. 1A**) were obtained as part of routine clinical practice to obtain specimens with diverse cell populations including intrinsic CNS cell populations such as microglia, neurons, and astrocytes, as well as immune populations that have originated from the periphery. These specimens were formalin-fixed and paraffin-embedded (FFPE) by the Northwestern University Neurological Surgery Tumor Bank.

**Fig. 1:**
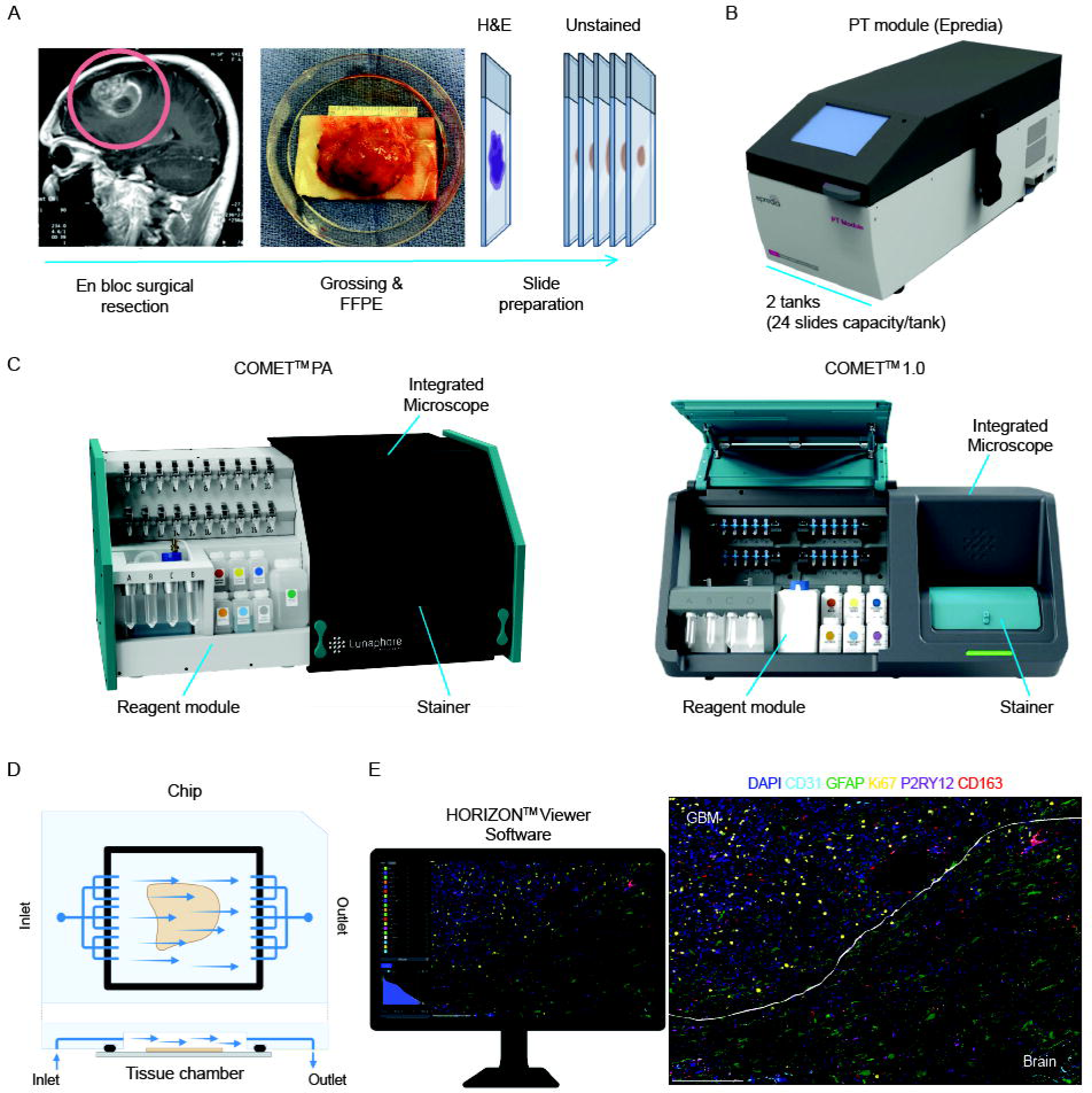
Workflow and introduction to the Lunaphore COMET™ benchtop system. **A**) The workflow of sample acquisition and preparation from the operating room to the lab. **B**) The Epredia^TM^ Lab Vision^TM^ PT module is used for the FFPE tissue slide dewaxing and antigen retrieval. **C**) Shown are images of the pre-commercial unit used in the study and the commercial Version 1.0 unit. The COMET^TM^ is composed of a reagent module (left compartment) and a staining and imaging module (right compartment). The docking area for 20x 2ml tubes represents the antibody reservoirs. After the completion of a 40-plex panel, these can be reloaded with another set of antibodies. The reagent module includes bottles and reservoirs for waste, reagents, and buffers. The staining and imaging module includes an integrated microscope with a 20x objective (not seen) and houses the rotating stage with 4 stainers. The COMET^TM^ control software and image viewing software are run with a Titan W599 Octane – Dual Intel Xeon Scalable CPU – Quad GPU CUDA Render Workstation PC. **D**) Illustration of the microfluidic chip [35]. The single-use chip that under pressure forms a closed chamber filled with buffer submerging the tissue sample and is used for staining and imaging acquisition. The maximum staining area is 9.1 x 9.1 mm. **E**) Representative view of the HORIZON™ Viewer software (version 0.1) with an example of a final generated tiff. snapshot (scale bar 200µm).

### Staining and Imaging Strategy

FFPE tissue slides were prepped (i.e., dewaxing and antigen retrieval) using the PT module (**Fig. 1B**) (Epredia) for 1 hour at 102°C in a Hier and Dewax Buffer solution pH=9 (Lunaphore) and run on the COMET™ system in which recurrent cycles of staining, imaging, and elution were performed (**Fig. 1C**). Two primary antibodies were stained at a time per cycle using the microfluidic chip (**Fig. 1D**), detected based on dichotomized immunofluorescence and subsequently imaged (**Fig. 1E**). Elution steps were conducted between cycles, to remove all primary and secondary antibodies before the start of a new cycle where two new primary antibodies are layered on the tissue followed by secondary antibodies. The complete removal of the antibodies during each elution step was confirmed using the optimization protocol that allows additional image acquisition and visualization after each elution cycle. Over the evaluation of 200+ antibodies at or below the manufacturer-recommended dilution (MRD), we did not identify any residual antibody staining after elution. We could label up to 40 markers per slide with ease only limited by the system reagent capacity. The COMET™ software automatically generates an OME-TIFF file combining all staining images for subsequent analysis. The quality control and the image visualization were performed using the HORIZON™ Viewer software (Lunaphore), which included the analysis of analyte location (nuclear, cytoplasmic, or membrane), analyte correlation or anti-correlation with other analytes, intensity, and signal-to-noise ratio; verified by a board-certified neuropathologist (C.H.). The OME-TIFF images post-processing, tissue and cell segmentation, phenotyping, and feature extraction were obtained using standardized commercial “Academic essential and Phenotype research edition” packages of the Visiopharm® software (currently version 9.2024).

### Autofluorescence dynamics and corrections

Automated seqIF™ begins with a quenching step to decrease tissue intrinsic autofluorescence followed by DAPI and the secondary channels acquisitions representing the first step in all the protocols available within the software (Optimization and SeqIF™). Baseline signals of autofluorescence can then be subtracted from biomarker images during the analysis step in the Viewer. For tissues in which autofluorescence remains an issue, UV light can be used by placing the slides in an unshielded window for exposure for a couple of hours before staining. Red blood cells (RBCs) are the most common culprit for autofluorescence in brain tissues which preferentially affects the 555/TRITC channel more than the 647/Cy5 channel. This confounder can be further negated during the downstream analysis of cell segmentation and with various analysis strategies [36]. Since the RBC lacks a nucleus, there is no associated DAPI staining. Cell identity is therefore based on the presence of the DAPI signal. Autofluorescence does not typically meet the percent coverage (quantile) above the threshold based on the background subtraction that occurs when the auto-fluorescent information is collected.

### Antibody optimization

Antibody validation and inclusion in a panel were performed using two major protocols: Optimization and Positioning, before finally being executed through a SeqIF™ protocol. The Optimization protocol is used for determining an appropriate primary antibody concentration. The Positioning protocol clarifies if the antibody needs to be placed at a specific cycle within the final SeqIF™ panel. Certain analytes perform better on the 647 channel which has a better signal-to-noise ratio compared to the 555 channel. In all protocols, the system stains and captures DAPI at every cycle which is used for the alignment and generation of the assembled image at the end of the protocol. After the background autofluorescence capture at the beginning of a protocol, the primary and secondary antibodies are applied with DAPI to create the first staining image during SeqIF^TM^. Thereafter, there are automated cycles of sequential elution, quenching, primary and secondary antibody staining along with DAPI, followed by imaging. Previous analysis, along with the current assessments of this strategy, indicates that elution is complete with no remaining signal on the tissue of either the primary or secondary antibodies [30]. The optimal antibody concentration chosen for each marker corresponds to the signal that most accurately fills the dynamic range of the camera at a given exposure time. When certain antibodies had a high signal-to-noise ratio, the concentration was further decreased, making the runs more cost-effective. Effectively, three different dilutions were tested for each primary antibody using the Optimization protocol, starting at MRD, then MRD/2, and MRD/4, and visually comparing the dynamic range signals generated at a fixed exposure time. After this step, a determination is made if a particular antibody cycle placement is needed using the Positioning protocol, which allows staining cycles after each 5 elution steps up to 20 total cycles emulating a full protocol for testing purposes of the stability of the epitope in between washes.

### Microglia marker

Most intrinsic cell populations within the CNS have well-established markers except microglia. TMEM119 is used to distinguish resident microglia from blood-derived macrophages in human brain tissues [37]. When protein expression data were evaluated using the Protein Atlas (www.proteinatlas.org), other immune cells such as B cells were noted to have expression. Staining of the human tonsil demonstrated positive dual expression of TMEM119 on CD20+ B cells and CD11c+ antigen-presenting cells (**Fig. 2A**). In contrast, the purinergic receptor P2RY12 was noted to be more specific to the microglia lineage using the Protein Atlas. Since there was no positive signal of P2RY12 in the tonsil, this marker was selected to demarcate microglia within the CNS. Similarly, glial fibrillary acidic protein (GFAP) stains astrocytes and most glioma tumor cells [38, 39]. This marker is typically included for most staining purposes even for non-glial tumors such as metastasis to identify the tumor-brain interface. For the adjacent relatively normal brain or non-tumor diseases, the brain architecture can be generally constructed using GFAP, MAP2-labeled neurons, and P2RY12 microglia (**Fig. 2B; Video 1**). Human tissues are not perfused before fixation and embedding. As such, intravascular circulating immune cells will be inadvertently included in the quantification and functional assessments of the tumor microenvironment (TME). The inclusion of a structural endothelial marker like CD31 is recommended when determining whether immune cells are within the TME. Within the Viewer, each marker is interrogated and assigned a pseudo-color by the operator to generate the Tiff. snapshots.

**Fig. 2:**
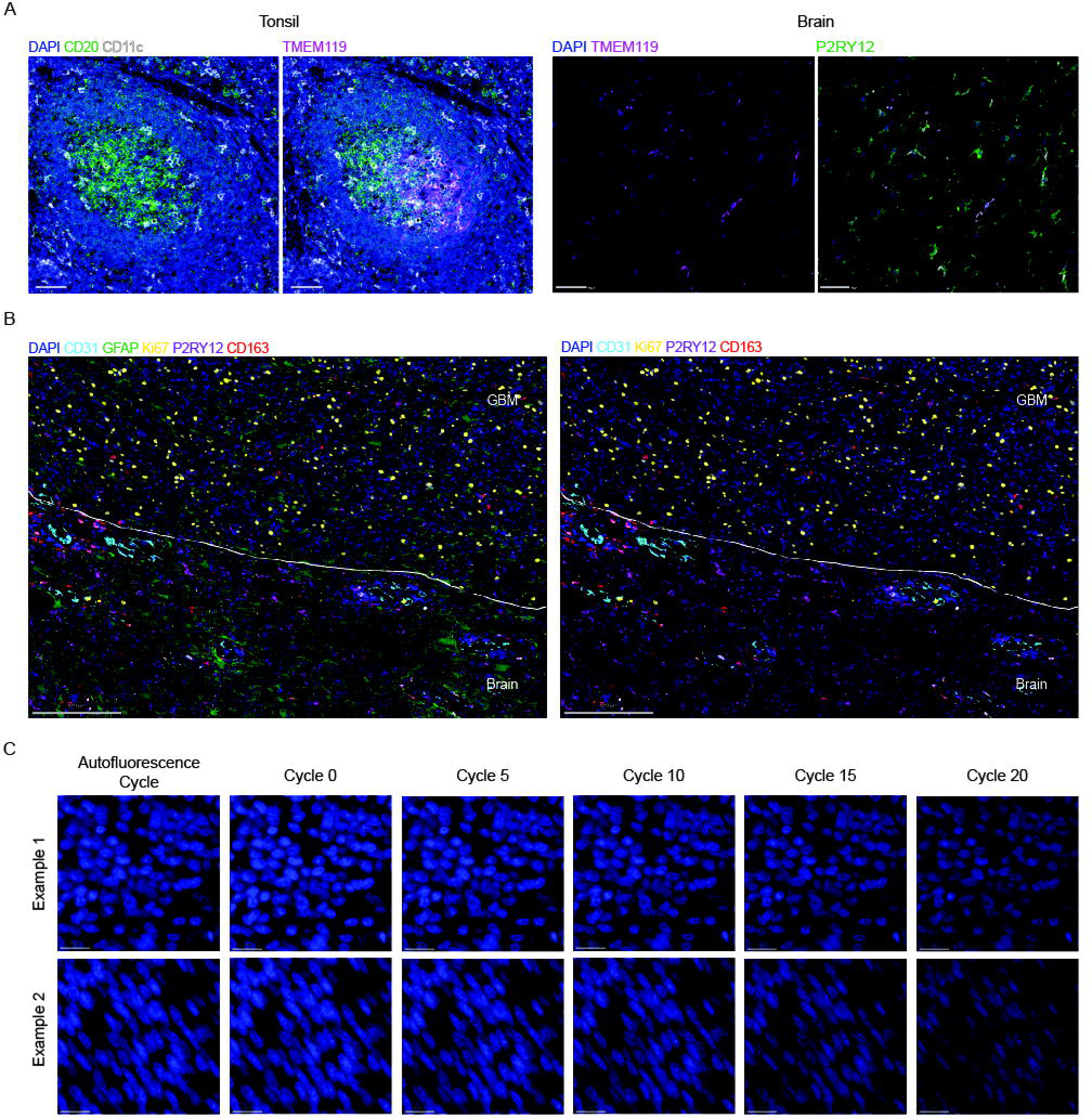
Cell lineage marker selection and nuclear capture. **A**) Selection of the microglia marker P2RY12 relative to TMEM119. Representative tonsil and brain (focal cortical dysplasia) multiplex IF images of the following markers: CD20 (B cells), CD11c (antigen-presenting cells), P2RY12 (microglia), and TMEM119 (microglia). Tonsil scale bars 100µm; Brain scale bars 50µm. **B**) Representative example of glioblastoma and adjacent brain and analyzed with the COMET™ benchtop system. The panel included markers for CD31 endothelium, GFAP astrocytes and glioma tumor cells, Ki67 proliferating tumor cells, P2RY12 microglia in the adjacent brain, and CD163 macrophages throughout the TME. **C**) Representative DAPI stained nuclei showing signal and nuclear morphology stability over elution cycles 0, 5, 10, 15, and 20. Example 1 is with background subtraction (BS). Example 2 is without BS during analysis. Scale bars 50µm.

### Nuclear morphology analysis

Because of the highly pleomorphic nature of the nuclei of glioblastoma, these specimens were used as representative examples. Using DAPI, the nuclei morphology in both adjacent normal brain and within glioblastoma specimens was captured after 0 and 20 elution cycles using an enabled image extraction feature after each cycle. There is no difference in the nuclear morphology but a slight decrease in the intensity of the DAPI staining was noted across cycles. This is accounted for by the COMET™ software algorithms while merging and stacking all images at the end of a protocol. The DAPI signal remains well above the detection threshold over 20 cycles (**Fig. 2C**).

### Antibody positioning and background subtraction assessments

Because the Lunaphore system involves sequential washes, there may be variability in epitope stability with subsequent cycles. Antibodies with targeted antigens resistant to this process should be positioned at later cycles. To ascertain if the antibodies should be run in a particular order and if the cellular compartment was associated with signal stability, nine targets were selected (**Suppl. Table 2**) and analyzed on human glioblastoma. These targets included cell lineage markers such as CD68 (monocytes), CD11c (antigen-presenting cells), GFAP (astrocytes), and P2RY12 (microglia), with the latter two cell types having fine ramified branches; architectural markers such as CD31 that identified the endothelial cells and ACTA2 for smooth muscle in the vascular wall selected for their usefulness in tissue orientation and analysis of immune cell distribution with the TME; two nuclear markers such as Ki-67 and p-STAT3; and the membrane-expressed marker HLA-DR. Each signal was visualized and quantitatively analyzed comparing the antibody staining pattern and mean intensities acquired within the images after one cycle of elution and staining versus 20 cycles. Most of the markers’ signals were identical regardless of the cycle. There was no change in cellular morphology between cycles for any marker. The signal intensity of Ki-67 and CD68 demonstrated a slight loss after the first 5 cycles of elution (Pearson correlation coefficient of 0.86 and 0.63 respectively) (**Fig. 3A, B**; **Suppl. Figs. 1-10**) when using the same threshold for all images and applying the background subtraction feature in the Viewer. Visual differences, especially with nuclear p-STAT3, are eliminated when the subtraction feature is not applied (Pearson coefficient of 0.96) (**Fig. 4A, B; Supplementary Figs. 1-10**). This indicates that there is no actual loss of the signal with washes but rather the signal is masked due to an overestimation of the background between cycle 1 and the subsequent cycles. As the autofluorescence cycle used for subtraction is the first cycle within a protocol, the background is at its highest intensity – especially for CNS tissues. In such scenarios, an alternative autofluorescence cycle can be generated after cycle 1 and used for subtraction during the image analysis. Most target staining signal intensity is the same regardless of the number of elution cycles and regardless of background subtraction during analysis (**Supplementary Figs. 11-14**). In circumstances of diminished signal intensity, these antibodies should be positioned in the first five cycles. Some markers however have a binary (i.e., on/off) expression pattern and show better signals in later cycles due to decreased background. Assigning markers to categories such as structural, phenotypic/lineage, and functional guides the overall selection for each custom project-oriented analysis (**Supplementary Table 2**).

**Fig. 3:**
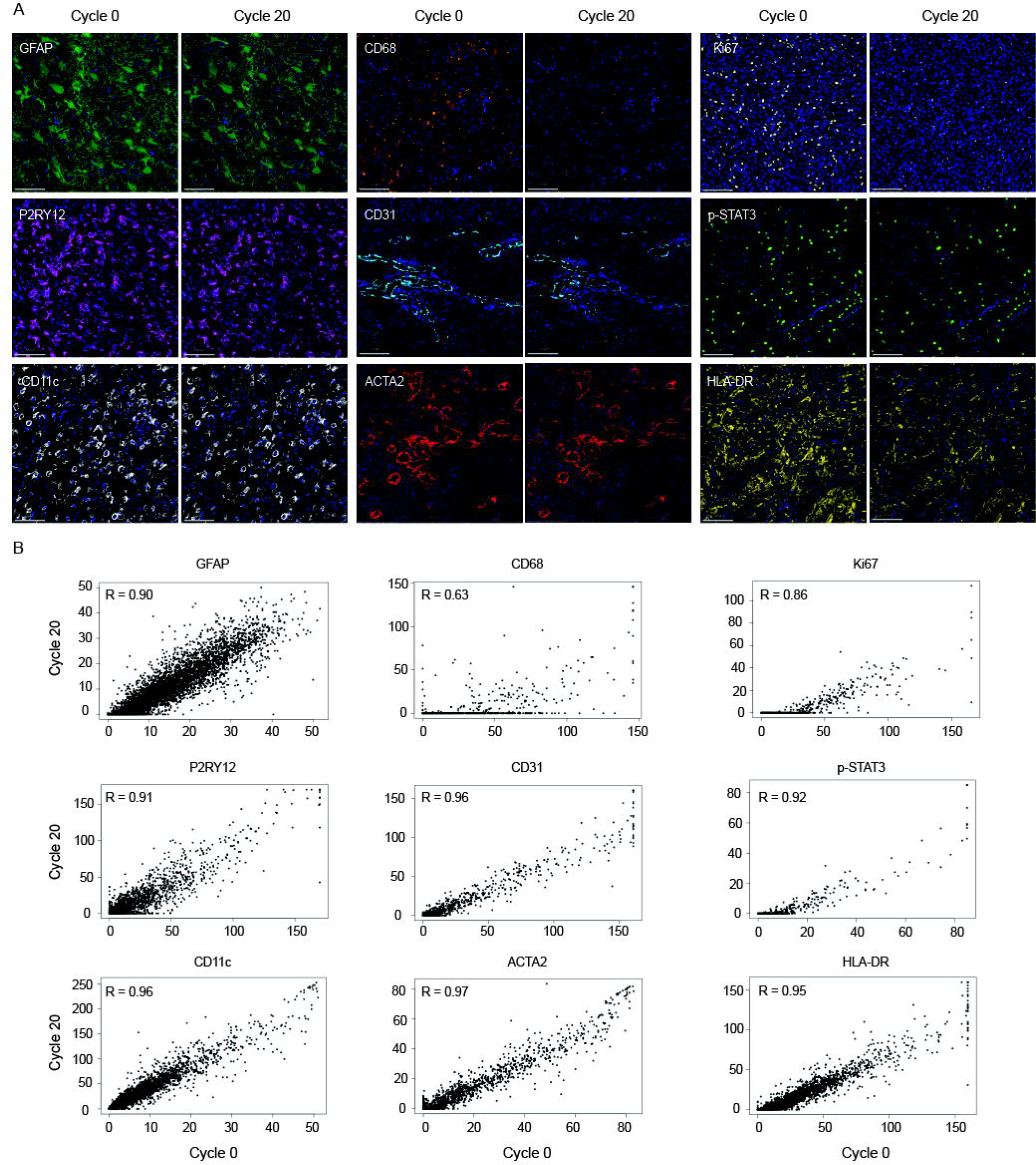
Marker imaging analysis with automated background subtraction (BS). **A**) Representative images, analyzed and captured with BS, of glioblastoma after elution cycles 0 and 20. The antibodies were directed to cell lineage and architectural markers GFAP, P2RY12, CD11c, CD68, vascular CD31 and ACTA2, functional nuclear markers Ki-67 and p-STAT3, and membrane HLA-DR. All scale bars at 100µm, except P2RY12 at 200µm. **B**) In Fiji, each image was converted to greyscale and then converted to a table, wherein each pixel was represented by a value between 0 (black) and 255 (white). Each image was then subtracted from the image after cycle 0 for that marker in the Fiji image calculator. For each cycle, 250 data points representing antibody pixels were randomly selected as individual data points. The data is represented using scatterplots showing the correlation between each antibody target intensity captured after cycles 0 and 20 with BS. In R, the Pearson correlation coefficient was calculated by comparing each pixel after cycle 0 to the same pixel after cycle 20 (all pixels, both antibody and background, were included in the analysis). Scatterplots were generated with 5000 randomly selected pixels (cf. Suppl. Fig. 2-10).

**Fig. 4:**
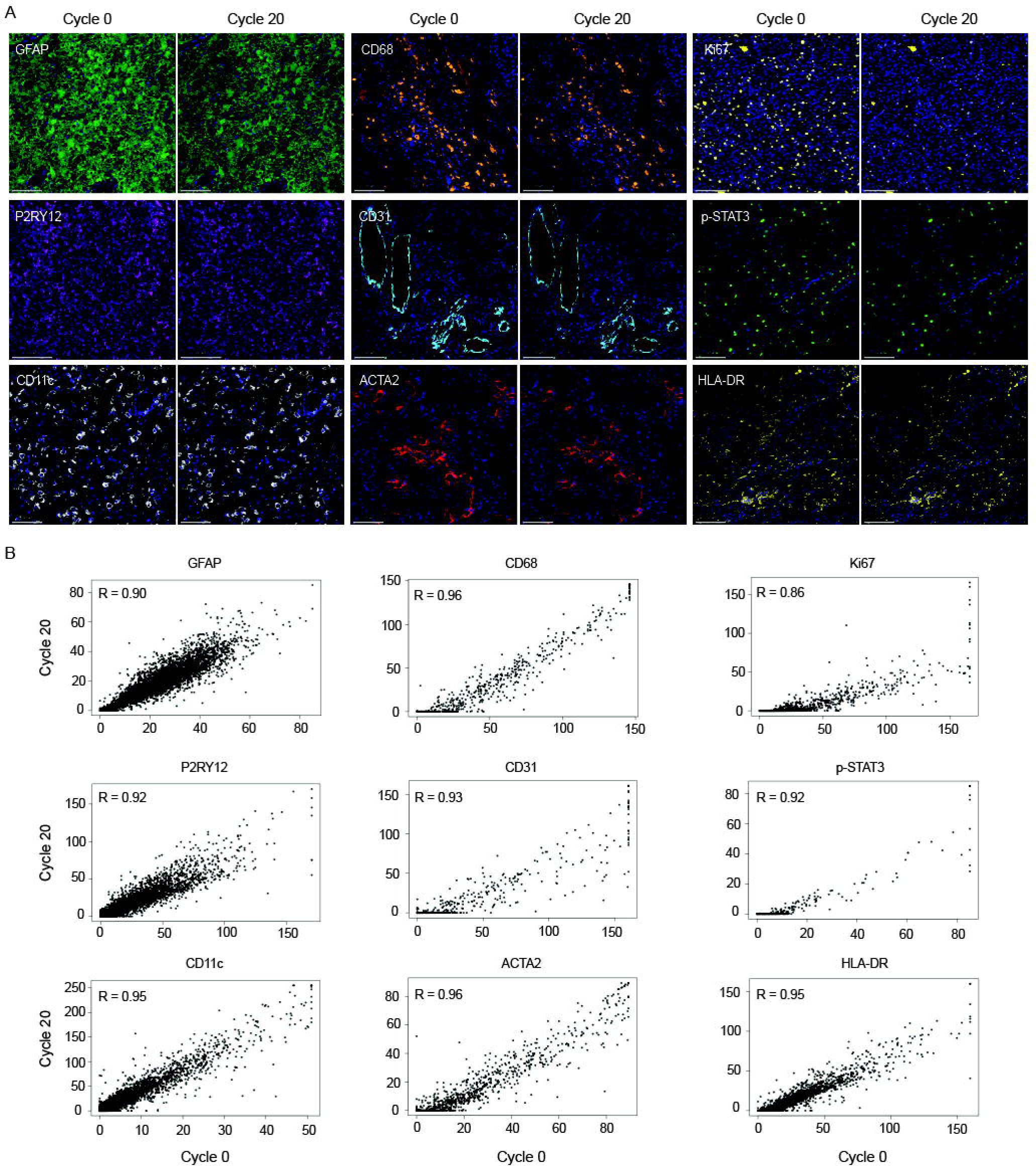
Marker imaging analysis without automated background subtraction (BS). **A**) Representative images, analyzed and captured without BS, of glioblastoma after elution cycles 0 and 20. The antibodies were directed to cell lineage and architectural markers GFAP, P2RY12, CD11c, and CD68, vascular CD31 and ACTA2, functional nuclear markers Ki-67 and p-STAT3, and membrane HLA-DR. All scale bars at 100µm, except P2RY12 at 200µm. **B**) In Fiji, each image was converted to greyscale and then converted to a table, wherein each pixel was represented by a value between 0 (black) and 255 (white). For each cycle, 250 data points representing antibody pixels were randomly selected as individual data points. The data is represented using scatterplots showing the correlation between each antibody target intensity captured after cycles 0 and 20 without background subtraction. In R, the Pearson correlation coefficient was calculated by comparing each pixel after cycle 0 to the same pixel after cycle 20 (all pixels, both antibody and background, were included in the analysis). Scatterplots were generated with 5000 randomly selected pixels (cf. Suppl. Fig. 2-10).

### Tissue loss characterization

No significant tissue loss was observed over any of the elution rounds with >95% of cells remaining after 20 cycles (**Fig. 5A**). This observation is independent of the type of tissue analyzed since there was no difference in tissue integrity or the fraction of cells in either glioblastoma or adjacent normal brain after 20 cycles of staining. Additionally, the morphology of the cells such as astrocytes and microglia, which have fine cellular extensions, were not altered with subsequent cycles (**Fig. 5B**). In comparison to standard immune fluorescence, the edge effect is mostly absent using the COMET™ platform which would preclude artifactual positive signals when the entire tissue is scanned for analysis. The edge effect is due to tissue lifting during processing minimized with the COMET™ microfluidic chip that maintains constant pressure during staining. Other multiplex imaging strategies have shown tissue loss attributed to quenching [35, 40].

### Reproducibility and batch normalization

Following antibody optimization, we assessed the reproducibility of COMET™ by analyzing the same glioblastoma section using the same panel on different dates and between different operators. The difference in percent positive cells between each replicate was small (mean difference 1-3%) using the same threshold settings for each marker in between images, and no significant difference was observed between operators (p>0.05). These data demonstrate reproducibility between batches and signal stability for each of the antibodies.

## CONCLUSION

The closest related systems to the COMET™ would be systems utilizing antibodies without modification or “off the shelf.” Typically, these systems utilize photobleaching or chemical stripping to multiplex. Most often these are academic-derived systems such as IBEX or CycIF [16]. Commercially, the ZellKraftWerk’s system evolved into the Bruker Canopy System, which was developed over a decade ago with minimal adoption [19]. Photobleaching, a component of these methods, uses an intense UV light source, which has heat as a by-product that creates tissue damage and loss through expansion and contraction of the tissue. Furthermore, the antibodies are not removed so there can be confounding issues of protein interactions. Notably, IBEX typically requires fresh frozen tissue which is not widely available in many instances. IBEX and CycIF are also entirely manual systems, the automation needed to standardize has not been created as they are largely academic systems. Without automation and prolonged staining times for IBEX and CycIF, both methods are relatively slower compared to COMET™. Furthermore, the extent of photobleaching is affected by tissue type and analysis of some tissues such as the brain may not be feasible with the Canopy System. Canopy, while it has evolved from its roots, still images through plastic windows and features high dynamic range imaging, which is in essence image averaging. For functional markers, the average intensity collected as it is degrading through photobleaching is not a quantitative metric.

The COMET™ is a commercial system assessed and validated for multiplex imaging interrogating CNS tissues. We also developed standard operating procedures and analysis throughput. This system is sufficiently simple and easy to maintain it can be used in an investigator’s laboratory. The COMET™ system allows for rapid panel development, is less time-consuming relative to manual protocols, and has less edge effect which confounds analysis of the tissue. Edge effect is especially relevant in the scenarios of analysis of limited tissue such as the case for brain biopsies or tissue microarrays. The COMET™ can be used to interrogate and quantify the entire tissue area, including the edges, as opposed to mere regions of interest. Antibody panels can be rapidly tailored to the specific interests of the research investigator as opposed to standardized panels because of the ease of antibody verification and incorporation.

## METHODS

### Study approval and consent

All human tissue was collected under Northwestern IRB HIPPA-approved protocols (STU00214485 and STU00214576) with the donors’ approval and consent. All tissue blocks were prepared using standard formalin fixation methods at the Neurological Surgery Tumor Bank of Northwestern University.

### Tissue preparation and antigen retrieval

FFPE slides were prepared at 4 microns thickness, mounted on positively charged slides, and stored at room temperature (RT). Hydration and deparaffinization steps of the tissue, along with antigen retrieval were done simultaneously using the PT module (PTM). The PTM (Epredia) includes two independently controlled antigen-retrieval fluid tanks that can harbor two slide racks each and up to 48 slides in one cycle. Epredia Dewax and Hier Buffers (15X, 100ml), H (cat #TA-999-DHBH; pH9) or L (cat #TA-999-DHBL; pH6) are prepared with deionized water (DIW) at 1:15 dilution, placed in the tank and preheated to maintain solution temperature at 35 to 85°C. Two different antigen retrieval protocols can be conducted simultaneously in each tank. After deposition of the slide racks within the tank and the lid is closed, the pre-treatment cycle is run for a 1-2h period, with a slow controlled increase in temperature to the boiling point of the solution at the beginning and cooling of solution at the end of the cycle to 102°C and 65°C, respectively. After the run is finished, a triple-beep sounds, notify the end of the cycle, unlocking the lid, and allowing the user to open and handle the slides. These latter are then placed in polyethylene Coplin jars including a polyethylene slide rack and washed 3x by Multi-staining Buffer (MSB) diluted with DIW at 1:20 dilution (Lunaphore BU06; 20X) for a couple of seconds each time while rocking the slide rack in a linear motion up and down. Slides are kept hydrated in MSB at RT until mounted onto the stainers within the COMET™ where a microfluidic system chip with an imaging area of 9x9mm is positioned on top of the tissue and the stainer closed to seal the staining chamber.

### Reagents and antibodies preparation

The sequential immunofluorescence multiplex staining conducted by the COMET™ (Lunaphore) is automated and controlled by its control software. Reagents and antibody volumes were automatically calculated and generated by the COMET™ Control software (Lunaphore) according to the planned protocol. The reagents were then prepared and placed in their appropriate locations within the COMET™ module. The reagents include MSB or washing buffer, Elution Buffer (Lunaphore BU07-L; solution 2 (20X) diluted in solution 1 (1X) at 1:20 dilution), Imaging Buffer (Lunaphore BU09; solution 1 (40X) prepared by dissolving the 25g solute powder in 60ml DIW and stored at -20°C; solution 2 (1X) prepared by dilution of solvent (10X) in DIW at 1:10 dilution and stored at RT; before use 1ml of solution 1 is diluted in 39ml of solution 2), Quenching Buffer (Lunaphore BU08-L; solution 1 (10X) diluted in DIW at 1:10 dilution; solution 2 (10X) diluted in solution 1 (1X) at 1:10 dilution), and 70% alcohol. Primary and secondary (Alexa Fluor 555 and 647) antibodies were diluted in MSB according to their optimized dilution and concentration and stored at 4°C until placed in the COMET™ reagent module. When all reagents were in their appropriate locations and the slides mounted on the stainers, the corresponding protocols were launched. The machine can run 4 slides of 20-plex panels over 24-30h.

### Sequential immunofluorescence (SeqIF™) multiplex staining

The multiplex panels were project-driven and customized according to the investigators’ hypothesis and scientific questions. We validated up to 200 CNS markers and targets (**Supplementary Table 1**), spanning multiple CNS diseases: adult brain and spine tumors (glioblastoma, meningioma, chordoma), pediatric low- and high-grade gliomas, non-tumor diseases like Alzheimer’s, epilepsy and IgG4 disease. For optimal concentration, signal, and noise detection, all antibodies were tested at three different dilutions, starting with MRD, then MRD/2, and MRD/4. Secondary Alexa Fluor 555 and 647 (ThermoFisher Scientific) were used at 1/200 and 1/400 dilutions respectively, in conjunction with the spectral 4’,6-diamidino-2-pheynlindole (DAPI; ThermoFisher Scientific) counterstain. All antibodies were validated using conventional immunohistochemistry and/or immunofluorescence staining. The optimizations (Optimization and Positioning) and full runs (Sequential IF protocols) of the multiplex panels were executed using sequential IF methodology integrated within the COMET™ platform. The staining corresponds to following automated cycles of 2 antibodies’ staining at a time, followed by imaging, and elution. During this process, no sample manipulation is required.

### Fluorescent microscopy and image registration

Image assembly and registration are executed within the COMET™ Control software immediately after concluding the staining procedure. Effectively, tissues are automatically scanned using the COMET™ integrated fluorescent high-power field imaging microscope/scan at 20x magnification. The microscope captures the fluorescent signals (DAPI, TRITC, and Cy5) separately at the corresponding wavelength with preset exposure times. The automated protocols in COMET™ result in a multi-layer OME-TIFF file, where the imaging outputs from each cycle are stitched and aligned. COMET™ OME-TIFF contains the DAPI image, intrinsic tissue autofluorescence in TRITC and Cy5 channels, and a single fluorescent layer per cycle in the TRITC and Cy5 channels. No tissue realignment issues nor disruption of the unique fluorescent signatures of the different markers were detected in the quality control check of the images.

### Autofluorescence removal and image analysis via the HORIZON™ Viewer

Unstained tissue images are acquired by default at the beginning of each COMET™ protocol and can be subsequently used as a reference for tissue-intrinsic autofluorescence. These images are enclosed as independent layers in multi-layer OME-TIFF and can be subtracted by pixel-wise subtraction, where the pixel value is normalized to the exposure time of different layers within the Viewer software. This subtraction occurs on the fly and does not affect the raw data in the OME-TIFF file. The operation can be canceled at any time. If background subtracted images are required for downstream analysis, they can be exported as new OME-TIFF files by the export function of the Viewer.

### Quantitative data analysis

Each antibody (n=9) was visualized individually without DAPI at cycles 0, 5, 10, 15, and 20 and analyzed with and without Lunaphore background subtraction. Image subtraction was performed to visualize changes in pixel intensity across cycles. Each image was opened in Fiji [41] and its properties were verified. The canvas size was adjusted to remove the portion of the image that contained the scale bar. Each image was then subtracted from the cycle 1 image for that marker in Fiji’s image calculator. All images, both original images, and subtractions, were transformed into greyscale, and then into a quantitative results table, in which the pixel intensity is represented by a number between 0 (black) and 255 (white). Boxplots were created to quantify changes in antibody intensity across cycles. Four randomly selected images (two different antibodies, with and without Lunaphore background subtraction) were opened in MIPAR [42] and basic pixel intensity thresholds were applied to each image. It was visually determined that for each image, the optimal threshold to segment the image background without also segmenting the borders of regions of antibody, was 10 units of intensity. The tables created by quantifying pixel intensity in Fiji were then read into R [43]. To focus only on pixels representing the antibody, data points representing pixels with an intensity less than or equal to 10 were removed from further calculation as these represent the background. Boxplots were created with the remaining data points for each cycle. Then, for each cycle, 250 data points representing antibody pixels were randomly selected and overlaid on the boxplot for that cycle as individual data points. For each antibody, the correlation was used to assess the antibody signal stability across cycles. The tables created by quantifying pixel intensities in Fiji were read into R and the Pearson correlation coefficient was calculated by comparing each pixel after cycle 0 to the same pixel after cycle 20 (all pixels, both antibody and background, were included in the analysis). A scatterplot was generated with 5000 randomly selected pixels, comparing their intensities after cycles 0 and 20. In R, data points less than or equal to 10 were removed from further calculation; then, for every antibody at each cycle, the mean pixel intensity and SEM were calculated. Line graphs of changes in antibody intensity across cycles were created in GraphPad Prism version 9.5.1 for Windows, GraphPad Software, Boston, Massachusetts USA, www.graphpad.com.

### Statistics and reproducibility

For each marker, a two-sample Wilcoxon rank-sum (unpaired) exact test was performed to compare the different conditions, and p-values were computed. To account for multiple comparisons, p-values were adjusted using the False Discovery Rate [44]. Pearson correlation analysis and coefficient were generated to compare the different techniques with ≥0.8 considered a strong correlation [30].

## SUPPLEMENTARY FIGURES AND VIDEOS

**Supplementary Figure 1: Summary graphs of mean antibody intensities across staining cycles 0, 5, 10, 15, and 20.** Mean antibody intensities across staining cycles with (**A**) or without (**B**) background subtraction. Error bars represent the standard error of the mean.

**Supplementary Figures 2-10: Visualization and analysis of markers’ signal intensity across cycles.** Fig. 2-10 representing CD68, GFAP, HLA-DR, P2RY12, ACTA2, p-STAT3, CD11c, Ki67, and CD31, respectively. Boxplots of antibody signal intensity with (**A**) or without (**B**) background subtraction (BS) across cycles. In Fiji, each image was converted to greyscale and then converted to a table, wherein each pixel was represented by a value between 0 (black) and 255 (white). To focus only on pixels representing antibodies, data points representing pixels with an intensity less than or equal to 10 (value determined by thresholding in MIPAR) were removed from the calculation as these represent the background. Boxplots were created with the remaining data points for each cycle. Then, for each cycle, 250 data points representing antibody pixels were randomly selected and overlaid on the boxplot for that cycle as individual data points. Images and image subtractions of antibody intensity with (**C**) or without (**E**) BS across cycles. After being captured by the COMET™ system, each image had its canvas size adjusted in Fiji to remove the portion of the image that contained the scale bar. Each image was then subtracted from the image after cycle 0 for that marker in the Fiji image calculator. Scatterplots showing a correlation between antibody intensities captured after cycles 0 and 20 with (**D**) and without (**F**) BS. In R, the Pearson correlation coefficient was calculated by comparing each pixel after cycle 0 to the same pixel after cycle 20 (all pixels, both antibody and background, were included in the analysis). Scatterplots were generated with 5000 randomly selected pixels. All scale bars at 50µm.

**Supplementary Fig. 11 and 12:** Series of multiplex IF images of different CNS markers captured after cycles 0, 5, 10, 15, and 20, and analyzed without background subtraction feature. The markers include (Fig. 11): CD68, Ki67, HLA-DR, CD11c, CD31, and (Fig. 12): GFAP, P2RY12, ACTA2, p-STAT3. All scale bars at 50µm.

**Supplementary Fig. 13 and 14:** Series of multiplex IF images of different CNS markers captured after cycles 0, 5, 10, 15, and 20, and analyzed with enabled background subtraction feature. The markers include (Fig. 13) CD68, Ki67, HLA-DR, CD11c, CD31, and (Fig. 14) GFAP, P2RY12, ACTA2, p-STAT3. All scale bars at 50µm.

**Video 1: A video representation of the brain architecture.** Multiplex images are shown sequentially in an assembled video of astrocytes (GFAP in orange), neurons (MAP2 in green), and microglia (P2RY12 in white) staining in normal brain adjacent to a region of resected focal cortical dysplasia.

## Acknowledgments

Special thanks to Laura Jaros for administrative support for the installation and setup of the Lunaphore Comet System.

## Funding

This work was supported by a gracious gift from the Stephen Coffman Trust and NIH grants CA120813, NS120547, and CA221747.

**Authors’ Contributions:**

Experimental design: HN, JT, ST, ABH

Software algorithms: DW

Samples preparation: HN, AS, KM, JW, CH

Funding acquisition: JC, RS, ML, ABH

Data analysis: HN, JT

Initial draft and writing: HN, JT, ABH, JB

Manuscript review: All authors approved the final manuscript.

**Conflict of interests: None**

