## Supplementary Tables for "High Dimensional Proteomic Multiplex Imaging of the Central Nervous System Using the COMET™ System"

**Supplementary Table 1: Validated primary and secondary antibodies**

| Marker | Company<br>Dilution | Host | Reactivity |
|---|---|---|---|
| **Primary antibodies** | | | |
| **GFAP** | 1* Abcam<br>Clone EPR1034Y<br>Ab68428<br>1/2000<br>2* Sigma<br>Clone GA5<br>MAB360<br>1/3000 | 1* Rabbit<br>2* Mouse | 1*Human<br>Rat<br>Mouse<br>2*Bovine<br>Chicken<br>Human<br>Mouse<br>Pig<br>Rat<br>Rabbit |
| **SOX2** | 1*Abcam<br>Ab92494<br>Clone EPR3131<br>1/5000<br>2*Abcam<br>Clone 20G5<br>Ab171380<br>1/1000 | 1* Rabbit<br>2* Mouse | 1*Human<br>2*Human<br>Mouse |
| **CD31** | Abcam<br>Clone EPR17259<br>Ab225883<br>1/1500 | Rabbit | Mouse<br>Rat<br>Human |
| **CD3** | Dako Agilent<br>Monoclonal F7.2.38<br>M7254<br>1/50 | Mouse | African green monkey<br>Hamster<br>Human<br>Mouse<br>Rat<br>Dog<br>Cow<br>Pig<br>Rabbit |

| | | | |
|---|---|---|---|
| **CD4** | 1* Abcam<br>Clone EPR6855<br>Ab133616<br>1/500<br>2* Abcam<br>Clone EPR19514<br>Ab183685<br>1/500 | Rabbit | 1* Human<br>2* Mouse |
| **CD8** | 1* Leica<br>Clone 4B11<br>PA0183<br>RTU<br>2* Cell signaling<br>Clone D4W2Z XP®<br>98941<br>1/500 | 1* Mouse<br>2* Rabbit | 1* Human<br>2* Mouse |
| **CD45RO** | 1*US Biological<br>Clone SPM125<br>172054<br>1ug/ml<br>2*Santa Cruz<br>Clone UCH-L1<br>Sc-1183<br>1/100 | Mouse | 1*Human<br>Chimpanzee<br>2*Mouse<br>Rat<br>Human |
| **CD19** | Abcam<br>Clone EPR5906<br>Ab134114<br>1/250 | Rabbit | Human |
| **CD20** | Dako Agilent<br>Clone L26<br>M0755<br>1/100 | Mouse | Human |
| **CD27** | Abcam<br>Clone EPR8569<br>Ab131254<br>1/1500 | Rabbit | Human |
| **CD138** | Dako Agilent<br>Clone MI15<br>M7228<br>1/100 | Mouse | Human |

| | | | |
|---|---|---|---|
| **CD11c** | 1* Abcam<br>Clone EP1347Y<br>Ab52632<br>1/300<br><br>2* Cell signaling<br>Clone D1V9Y<br>97585<br>1/300 | Rabbit | 1* Human<br><br>2* Mouse |
| **CD205** | Abcam<br>Clone EPR5233<br>ab124897<br>1/700 | Rabbit | Mouse<br>Human |
| **TMEM119** | 1*Abcam<br>Polyclonal<br>ab185333<br>1/150<br><br>2*Abcam<br>Clone 28-3<br>Ab209064<br>1/100 | Rabbit | 1*Human<br><br>2*Mouse |
| **NKG2D** | 1*Novus Bio<br>NB100-65956<br>Clone 1D11<br>1/500<br><br>2*Santa Cruz<br>Clone A7<br>Sc-515599<br>1/100 | Mouse | 1*Human<br><br>2*Mouse<br>Rat |
| **CD68** | 1* Dako Agilent<br>Clone PG-M1<br>GA613<br>RTU<br>Or M0876<br>Concentrate<br>1/200<br><br>2* Santa Cruz<br>Clone KP1<br>Sc-20060<br>1/200 | Mouse | 1* Human<br><br>2* Mouse |

| | | | |
|---|---|---|---|
| **CD163** | Abcam<br>Clone EPR19518<br>Ab182422<br>1/600 | Rabbit | Mouse<br>Rat<br>Human |
| **cFOS** | Abcam<br>Clone 2H2<br>ab208942<br>1/300 | Mouse | Mouse<br>Rat<br>Human |
| **MAP2** | Abcam<br>Clone EPR19691<br>Ab183830<br>1/100 (from stock at 1/1000) | Rabbit | Mouse<br>Rat<br>Human |
| **PSD95** | Abcam<br>ab18258<br>1/500 | Rabbit | Mouse<br>Rat<br>Human |
| **VGlut1** | Novus Biologicals<br>NBP2-46627<br>Clone CL2754<br>1/5000 | Mouse | Human<br>Mouse<br>Rat |
| **BIII tubulin** | Biotechne R&D systems<br>MAB1195<br>Clone TuJ-1<br><br>E: 1/100 (from stock at 1/100) | Mouse | Human<br>Mouse<br>Rat |
| **p-STAT3** | Cell Signaling Tech<br>(Tyr705) (D3A7) XP<br>9145S<br>1/250 | Rabbit | Human<br>Mouse<br>Rat<br>Monkey |
| **FOXP3** | 1*Cell signaling tech<br>Clone D2W8E<br>98377S<br>1/100<br><br>2*Abcam<br>EPR22102-37<br>Ab215206<br>1/100 | Rabbit | 1*Human<br>2*Mouse<br>Rat<br>Human |

| | | | |
|---|---|---|---|
| **p-ERK1/2** | Cell Signaling<br>D13.14.4E<br>4370<br>1/1700 | Rabbit | Human<br>Mouse<br>Rat<br>Hamster<br>Monkey<br>Mink<br>D.melanogaster<br>Zebrafish<br>Bovine<br>Dog<br>Pig<br>S. cerevisiae |
| **PD-1** | 1*Abcam<br>Clone EPR4877(2)<br>Ab137132<br>1/400<br>2*Abcam<br>EPR20665<br>Ab214421<br>1/500 | Rabbit | 1*Human<br>2*Mouse |
| **PD-L1** | GenomeMe<br>IHC411-1<br>Clone IHC411<br>1/100 | Rabbit | 1*Human<br>2*Mouse<br>Rat<br>Human |
| **IFN-γ** | Abcam<br>Ab231036<br>Clone EPR21704<br>1/500 | Rabbit | Human |
| **Perforin** | 1* Leica<br>Clone 5B10<br>NCL-L-Perforin<br>1/100<br>2* Santa Cruz<br>Clone F-1<br>Sc-136994<br>1/500 | Mouse | 1* Human<br>2* Mouse<br>Rat<br>Human |
| **LCK** | Cell signaling tech<br>Clone D88 XP<br>2984S<br>1/100 | Rabbit | Human |

| | | | |
|---|---|---|---|
| **p-LCK** | Novus Bio<br>MAB7500<br>Clone 755103 (p Tyr394)<br>1/500 | Mouse | Human |
| **LFA-1**<br>**(CD11a)** | 1* Abcam<br>Clone EP1285Y<br>Ab52895<br>1/1000<br><br>2* GeneTex<br>Clone 38<br>GTX75442<br>1/1000 | 1* Rabbit<br>2* Mouse | 1* Human<br>2* Mouse<br>Human |
| **ICAM-1**<br>**(CD54)** | 1*BioRad<br>Clone 15.2<br>MCA1615<br>1/200<br><br>2*Santa Cruz<br>G-5<br>Sc-8439 | Mouse | 1*Human<br>Pig<br><br>2*Human<br>Mouse<br>Rat |
| **CD66b** | Novus Biologicals<br>Clone G10F5<br>NB100-77808<br>1/50 | Mouse | Human<br>Mouse |
| **HLA I**<br>**(A, B, & C)** | Abcam<br>Clone EMR8-5<br>Ab70328<br>1/3000 | Mouse | Human |
| **HLA DR** | Abcam<br>Clone TAL 1B5<br>Ab20181<br>1/1100 | Mouse | Human |
| **MHC II** | Abcam<br>MRC OX-6<br>Ab23990<br>1/500 | Mouse | Mouse<br>Rat |
| **p-Chek2** | Cell signaling tech<br>Clone (Thr68) C13C1<br>2197S | Rabbit | Human |

| | | | |
|---|---|---|---|
| | 1/200 | | |
| **STING** | Cell signaling tech<br>Clone D2P2F<br>13647S<br>1/100 | Rabbit | Human<br>Mouse |
| **Podoplanin** | Leica<br>Clone D2-40<br>PA0796<br>RTU | Mouse | Human<br>Mouse<br>Rat |
| **Somatostatin Receptor 2 (SSTR2)** | Abcam<br>Clone UMB1<br>Ab134152<br>1/2000 | Rabbit | Human<br>Mouse<br>Rat |
| **Notch3** | Millipore Sigma<br>Clone 1E4<br>MABC594<br>1/50 | Rabbit | Human<br>Mouse |
| **CD73** | Cell signaling tech<br>Clone D7F9A<br>13160S<br>1/100 | Rabbit | Human<br>Mouse<br>Rat |
| **TIGIT** | Cell signaling tech<br>Clone E5Y1W<br>99567S<br>1/100 | Rabbit | Human |
| **TIM-3** | Abcam<br>Clone EPR22241<br>Ab241332<br>1/1000 | Rabbit | Human<br>Mouse<br>Rat |
| **LAG-3** | Novus Bio<br>Clone L4-PL33<br>(17B4)<br>NBP1-97657<br>1/500 | Mouse | Human |

| | | | |
|---|---|---|---|
| **IDO** | Invitrogen<br>Clone V1NC3IDO<br>14-9750-82<br>0.5µg/ml | Mouse | Human |
| **SIRP-α** | Cell signaling tech<br>Clone D613M<br>13379<br>1/100 | Rabbit | Human<br>Mouse<br>Rat<br>Monkey |
| **Granzyme B** | Santa Cruz<br>Clone 2C5<br>Sc-8022<br>1/500 | Mouse | Human<br>Mouse<br>Rat |
| **COL1A** | Abcam<br>EPR7785<br>Ab138492<br>1/1000 | Rabbit | Human |
| **COL4A** | Abcam<br>Polyclonal<br>Ab6586<br>1/100 | Rabbit | Human<br>Mouse<br>Rat<br>Hamster<br>Cow<br>Dog<br>Pig<br>Zebrafish<br>African green monkey<br>Chinese hamster<br>Syrian hamster |
| **TGF-β** | Abcam<br>Polyclonal<br>Ab92486<br>1/100 | Rabbit | Human |

| | | | |
|---|---|---|---|
| **BRAF V600E** | BioCare Medical (Ventana Medical Systems, Inc.)<br>790–4855<br>Clone VE1<br>RTU | Mouse | Human |
| **Brachyury** | Abcam<br>Polyclonal<br>Ab20680<br>1/100 | Rabbit | Mouse<br>Human |
| **NeuN** | 1*Millipore Sigma<br>Clone A60<br>MAB377<br>1/500<br><br>2*Cell signaling<br>Clone D4G4O XP®<br>24307<br>1/400 | 1*Mouse<br><br>2*Rabbit | 1*Avian<br>Chicken<br>Ferret<br>Human<br>Mouse<br>Pig<br>Rat<br>Salamander<br><br>2*Human<br>Mouse<br>Rabbit |
| **P2RY12** | 1* Atlas Antibodies<br>Polyclonal<br>HPA014518<br>1/1000<br><br>2* Cell signaling<br>Clone E9J1J<br>69766<br>1/1000 | Rabbit | 1* Human<br>2* Mouse |
| **CTLA-4 (CD152)** | Abcam<br>Clone CAL49<br>Ab237712<br>1/100 | Rabbit | Human<br>Mouse |
| **FC-γ R-IIIA (CD16A)** | Abcam<br>Clone SP175<br>Ab183354<br>1/100 | Rabbit | Human |

| | | | |
|---|---|---|---|
| **B7H3** | Cell signaling tech<br>Clone D9M2L<br>14058<br>1/100 | Rabbit | Human |
| **CD1e** | Abcam<br>Clone EPR15746(B)<br>Ab187157<br>1/100 | Rabbit | Human<br>Mouse<br>Rat |
| **ACTA2** | Abcam<br>Clone 1A4<br>Ab7517<br>1/1000 | Mouse | Human<br>Rat |
| **Vimentin** | Cell signaling<br>D21H3 XP<br>5741<br>1/700 | Rabbit | Human<br>Mouse<br>Rat<br>Monkey |
| **PDGFRA** | Cell signaling<br>D1E1E XP<br>3174<br>1/100 | Rabbit | Human<br>Mouse |
| **P16** | Roche<br>Clone E6H4<br>06680003001<br>RTU | Mouse | Human |
| **IL6Rα** | ThermoFisher<br>Polyclonal<br>PA5-102425<br>1/50 | Rabbit | Human, Mouse |
| **Synaptophysin** | Abcam<br>Clone SY38<br>Ab8049<br>1/500 | Mouse | Human<br>Mouse<br>Rat<br>Hamster<br>Cow |

| | | | |
|---|---|---|---|
| **Gal9** | Cell signaling Tech<br>Clone D9R4A<br>54330S<br>1/100 | Rabbit | Human |
| **SHH** | Abcam<br>Clone EP1190Y<br>Ab53281<br>1/1000 | Rabbit | Human |
| **IHH** | Abcam<br>Polyclonal<br>Ab39634<br>1/100 | Rabbit | Human<br>Mouse |
| **CD45** | 1* Dako Agilent<br>Clone 2B11 +<br>PD7/26<br>M070101-2<br>1/100<br><br>2* Abcam<br>Polyclonal<br>Ab10558<br>1/2000 | 1* Mouse<br>2* Rabbit | 1* Human<br>2* Mouse<br>Rat<br>Human |
| **p-TBK1** | Cell signaling<br>Clone D52C2<br>5483S<br>1/100 | Rabbit | Human<br>Mouse |
| **p-IRF3** | Cell signaling<br>Clone 4D4G<br>4947S<br>1/1500 | Rabbit | Human<br>Mouse |

| | | | |
|---|---|---|---|
| **iNOS** | Santa Cruz<br>Clone C-11<br>Sc-7271 (200µg/ml)<br>1/100 | Mouse | Human<br>Mouse<br>Rat |
| **PTPRZ1** | Atlas Antibodies<br>Polyclonal<br>HPA015103<br>1/700 | Rabbit | Human |
| **Ki67** | 1*DAKO Agilent<br>MIB-1 clone<br>GA62661-2<br>RTU<br><br>*2 BD Biosciences<br>Clone B56<br>550609<br>1/500<br><br>3* Abcam<br>Clone SP6<br>Ab16667<br>1/500 | 1-2* Mouse<br><br>3* Rabbit | 2-3* Human<br>Mouse<br>Rat<br><br>1*Human |
| **LYVE-1** | Abcam<br>Polyclonal<br>ab14917<br>1/100 | Rabbit | Mouse<br><br>Human |
| **INPP4B** | Abcam<br>EPR3108Y<br>Ab81269<br>1/50 | Rabbit | Human |
| **VISTA (B7-H5)** | Fortis Life Sciences<br>Clone BLR035F<br>A700-035<br>1/500 | Rabbit | Human |

| | | | |
|---|---|---|---|
| **pan-RAS** | ThermoFisher<br>(Invitrogen)<br>Clone Ras10<br>MA1-012<br>1/200 | Mouse | Human<br>Mouse |
| **TCRγδ** | ThermoFisher<br>(Invitrogen)<br>Clone 5A6.E9<br>TCR1061<br>1:200 | Mouse | Human |
| **CXCL16** | 1* LSBio<br>Polyclonal<br>LS-B8223<br>10ug/ml | 1* Rabbit | 1* Human |
| **CXCR6** | Abcam<br>Polyclonal<br>Ab8023<br>20µg/ml | Rabbit | Human |
| **β amyloid** | Biolegend<br>Clone 4G8<br>800701<br>1/1000 | Mouse | Human<br>Mouse |
| **Tau** | ThermoFisher<br>(Invitrogen)<br>AT8<br>MN1020<br>1/1500 | Mouse | Human<br>Mouse<br>Rat |
| **CD34** | Abcam<br>Clone EP373Y<br>Ab81289<br>1/100 | Rabbit | Mouse<br>Rat<br>Human |

| | | | |
|---|---|---|---|
| **Olig2** | Abcam<br>Clone EPR2673<br>Ab109186<br>1/100 | Rabbit | Mouse<br>Human<br>Rat |
| **Prohibitin** | Abcam<br>Clone EP2803Y<br>Ab75766<br>1/200 | Rabbit | Human<br>Mouse<br>Rat |
| **GAB1** | Cell signaling<br>Polyclonal<br>3232<br>1/100 | Rabbit | Human<br>Mouse<br>Rat<br>Monkey |
| **FAAH** | Abcam<br>Clone 4H8<br>Ab54615<br>1/100 | Mouse | Human<br>Mouse<br>Rat |
| **GPR183** | My Bio Source<br>Polyclonal<br>MBS7000994<br>1/500 | Rabbit | Human |
| **LGMN (Legumain)** | Cell signaling<br>Clone D6S4H<br>93627<br>1/1000 | Rabbit | Human<br>Mouse<br>Rat |
| **CD206** | Abcam<br>Polyclonal<br>Ab64693<br>1/1000 | Rabbit | Human<br>Mouse<br>Rat |
| **CD40** | Abcam<br>Polyclonal<br>Ab13545<br>1/500 | Rabbit | Human<br>Mouse<br>Rat<br>Sheep<br>Rabbit<br>Guinea pig |

| | | | |
|---|---|---|---|
| | | | Hamster<br>Cow<br>Pig<br>Monkey |
| **CD47** | Novus Bio<br>Clone B6H12.2<br>NBP2-31106<br>1/100 | Mouse | Human<br>Mouse |
| **DBI** | Invitrogen<br>Polyclonal<br>PA5-89139<br>1/500 | Rabbit | Human<br>Mouse<br>Rat |
| **CX3CR1** | 1*Biolegend<br>Clone 8E10.D9<br>824001<br>1/100<br><br>2*ThermoFisher<br>Clone 1H14L7<br>702321<br>1/100 | 1* Mouse<br>2* Rabbit | 1* Human<br>2* Human<br>Mouse |
| **GABA** | Sigma<br>Polyclonal<br>A2052<br>1/350 | Rabbit | Wide range |
| **GABARAP** | Abcam<br>EPR4805<br>Ab109364<br>1/600 | Rabbit | Human<br>Mouse<br>Rat |
| **GPNMB** | 1* Cell signaling<br>E4D7P XP<br>38313B<br>1/100<br><br>2* ProteinTech<br>66926-1-Ig<br>Clone 2B10B8<br>1/500 | 1*Rabbit<br>2*Mouse | 1*Human<br>2*Human<br>Mouse<br>Rat |

| | | | |
|---|---|---|---|
| **c-Caspase 3** | Cell signaling<br>Asp175<br>9661S<br>1/250 | Rabbit | Human<br>Mouse<br>Rat<br>Monkey |
| **NFAT1 (NFATc2)** | Abcam<br>Ab2722<br>Clone 25A10.D6.D2<br>1/500 | Mouse | Human |
| **NFAT2 (NFATc1)** | Santa Cruz<br>Sc-7294<br>Clone 7A6<br>1/500 | Mouse | Human<br>Mouse<br>Rat |
| **COL6A** | 1* Abcam<br>EPR17072<br>Ab182744<br>1/1000<br><br>2* Sigma<br>HPA010080<br>Polyclonal<br>1/100 | Rabbit | 1*Human<br>Mouse<br>Rat<br><br>2*Human |
| **LAIR-1** | 1* Santa Cruz<br>Clone F-5<br>Sc-398141<br>1/100<br><br>2* Sigma<br>HPA011155<br>Polyclonal<br>1/500 | 1*Mouse<br><br>2*Rabbit | Human<br>Mouse<br>Rat |
| **PDGFRB** | Cell signaling tech<br>Clone 28E1<br>3169<br>1/100 | Rabbit | Human<br>Mouse<br>Rat |

| | | | |
|---|---|---|---|
| **LOXL2** | Abcam<br>Ab96233<br>Polyclonal<br>1/100 | Rabbit | Human<br>Mouse |
| **TNF-α** | Abcam<br>Clone 52B83<br>Ab1793<br>1/100 | Mouse | Human<br>Mouse<br>Guinea pig<br>Chimpanzee<br>Zebrafish<br>Cynomolgus monkey<br>Rhesus monkey<br>Apteronotus leptorhynchus |
| **GALR-3** | LSBio<br>Polyclonal<br>LS-A204-50<br>1/500 | Rabbit | Human<br>Mouse<br>Rat<br>Dog |
| **CD71** | ThermoFisher<br>Clone H68.4<br>13-6800<br>1/200 | Mouse | Human<br>Mouse<br>Rat<br>Hamster<br>Chicken |
| **HIF-1α** | Novus Bio<br>NB100-105<br>Clone H1alpha67<br>1/200 | Mouse | Human<br>Mouse<br>Rat<br>Canine<br>Bovine<br>Porcine<br>Primate<br>Rabbit<br>Sheep<br>Feline<br>Mammal<br>Xenopus |
| **Cytokeratin** | Dako Agilent<br>M3515 529-2<br>Clone AE1/AE3<br>1/100 | Mouse | Human |

| | | | |
|---|---|---|---|
| **Gelsolin** | Cell signaling<br>Clone D9W8Y<br>12953<br>1/600 | Rabbit | Human<br>Mouse<br>Rat<br>Monkey |
| **Versican (VCAN)** | Sigma<br>HPA004726<br>Polyclonal<br>1/500 | Rabbit | Human |
| **Fibrinogen** | Abcam<br>Ab34269<br>Polyclonal<br>1/500 | Rabbit | Mouse<br>Rabbit<br>Human |
| **Fibronectin** | Abcam<br>Ab2413<br>Polyclonal<br>1/1000 | Rabbit | Human<br>Mouse |
| **Myo1C** | Sigma<br>HPA001768<br>Polyclonal<br>1/200 | Rabbit | Human |
| **CD204 (MRS1)** | Affinity Biosciences<br>Polyclonal<br>DF6694<br>1/1000 | Rabbit | Human<br>Mouse<br>Rat |
| **EMA (MUC-1)** | Abcam<br>Polyclonal<br>Ab15481<br>1/1000 | Rabbit | Human<br>Mouse |
| **CD44** | Abcam<br>Clone EPR18668<br>Ab189524<br>1/5000 | Rabbit | Human<br>Mouse<br>Rat |

| | | | |
|---|---|---|---|
| **VEGFA** | Abcam<br>Clone VG-1<br>Ab1316<br>1/1000 | Mouse | Human<br>Mouse<br>Rat |
| **HES1** | Cell signaling<br>D6P2U<br>11988<br>1/2000 | Rabbit | Human<br>Mouse<br>Rat<br>Monkey |
| **TTYH1** | Novus Bio<br>Polyclonal<br>NBP1-59909<br>1/500 | Rabbit | Human<br>Mouse |
| **CENPF** | Abcam<br>Polyclonal<br>Ab5<br>1/750 | Rabbit | Human<br>Mouse |
| **DYNLRB2** | ThermoFisher<br>Polyclonal<br>PA5-23763<br>1/100 | Rabbit | Human<br>Mouse |
| **ZIC1** | Novus Bio<br>Polyclonal<br>NB600-488<br>1/500 | Rabbit | Human<br>Mouse<br>Rat |
| **PBX1** | Cell signaling<br>Polyclonal<br>4342S<br>1/400 | Rabbit | Human<br>Mouse |
| **PAX6** | Abcam<br>Polyclonal<br>Ab5790<br>1/200 | Rabbit | Human<br>Mouse<br>Rat<br>Monkey |

| | | | |
|---|---|---|---|
| **NFIB** | Abcam<br>EPR14122<br>Ab186738<br>1/500 | Rabbit | Human<br>Mouse<br>Rat |
| **FABP7** | ThermoFisher<br>Polyclonal<br>PA5-24949<br>1/500 | Rabbit | Human<br>Mouse |
| **Ascl1** | Abcam<br>EPR19840<br>Ab211327<br>1/100 | Rabbit | Human<br>Mouse |
| **Gdf10** | GeneTex<br>Clone N3C3<br>GTX118039<br>1/1000 | Rabbit | Human<br>Mouse<br>Rat |
| **PTPRZ** | Abcam<br>Polyclonal<br>Ab126497<br>1/500 | Rabbit | Human |
| **ATP1A2** | Abcam<br>M7-PB-E9<br>Ab2871<br>1/500 | Mouse | Human<br>Mouse<br>Rat |
| **SOX4** | ThermoFisher<br>Clone CL5665<br>MA5-31424<br>1/2000 | Mouse | Human<br>Mouse<br>Rat |
| **PAX3** | Biotechne<br>Clone 274212<br>MAB2457<br>1/1000 | Mouse | Human<br>Mouse |

| | | | |
|---|---|---|---|
| **DACH1** | Proteintech<br>Clone 3B6D2<br>60082-1-Ig<br>1/500 | Mouse | Human<br>Mouse<br>Rat |
| **BOC** | Biotechne<br>Clone 273729<br>MAB20361<br>1/100 | Mouse | Human<br>Mouse |
| **Nestin** | 1* Sigma<br>Clone rat-401<br>MAB353<br>1/200<br><br>2* Sigma<br>Clone 10C2<br>MAB5326<br>1/200 | Mouse | 1* Mouse<br>Rat<br><br>2* Human |
| **WLS** | ThermoFisher<br>Clone A7C2<br>MA5-44978<br>1/500 | Mouse | Human<br>Mouse<br>Rat |
| **Aqp4** | Abcam<br>Clone 4/18<br>Ab9512<br>1/200 | Mouse | Human<br>Mouse<br>Rat<br>Rabbit<br>Zebrafish |
| **Msx1** | Novus Bio<br>Clone 1E2<br>H00004487-M11<br>1/100 | Mouse | Human<br>Mouse |
| **Foxj1** | ThermoFisher<br>Clone 2A5<br>14-9965-80<br>1/1000 | Mouse | Human<br>Mouse<br>Pig<br>Rat<br>Fruit Fly |
| **OLIG1** | Sigma<br>Monoclonal<br>MAB5540<br>1/500 | Mouse | Human<br>Mouse<br>Rat |

| | | | |
|---|---|---|---|
| **ApoE** | Novus Bio<br>Clone WUE-4<br>NB110-60531<br>1/500 | Mouse | Human<br>Mouse |
| **CRABP1** | Santa Cruz<br>Clone F-9<br>Sc-166897<br>1/500 | Mouse | Human<br>Mouse<br>Rat |
| **CD103** | Abcam<br>EPR4166(2)<br>Ab129202<br>1/500 | Rabbit | Human |
| **SLFN11** | Cell signaling<br>D8W1B<br>34858S<br>1/200 | Rabbit | Human |
| **SLFN5** | Sigma<br>Polyclonal<br>HPA017760<br>1/200 | Rabbit | Human |
| **ROR-γ** | Santa Cruz<br>Clone D-4<br>Sc-365476<br>1/100 | Mouse | Human |
| **AR** | Abcam<br>EPR1535(2)<br>Ab133273<br>1/300 | Rabbit | Human<br>Mouse<br>Rat |
| **CD101 (IGSF2)** | Proteintech<br>Polyclonal<br>26047-1-AP<br>1/500 | Rabbit | Human<br>Mouse |
| **GLUT5 (SLC2A5)** | ThermoFisher<br>(Invitrogen)<br>MA1-036X<br>Clone 14C8<br>1/100 | Mouse | Human<br>Mouse |

| | | | |
|---|---|---|---|
| **CCR2** | Novus Bio<br>NBP1-48338<br>Polyclonal<br>1/200 | Rabbit | Human<br>Mouse |
| **JAM-A** | ThermoFisher<br>(Invitrogen)<br>Polyclonal<br>36-1700<br>1/500 | Rabbit | Human<br>Mouse<br>Rat<br>Dog<br>Hamster |
| **IDH1 Mut** | Dianova<br>DIA-H09-L<br>Clone H09<br>RTU | Mouse | Human |
| **TERT** | Novus Bio<br>NB100-317<br>Clone 2C4<br>1/100 | Mouse | Human<br>Mouse<br>Rat |
| **Yap** | Santa Cruz<br>Sc-101199<br>Clone 63.7<br>1/1000 | Mouse | Human<br>Mouse<br>Rat |
| **Taz** | Cell Signaling<br>4883S<br>Clone V386<br>1/1000 | Rabbit | Human<br>Mouse<br>Rat |
| **CD5** | Cell Signaling<br>39300<br>Clone E8X3S XP®<br>1/800 | Rabbit | Human<br>Monkey |
| **CCR7** | Biotechne<br>MAB197<br>Clone 150503<br>1/100 | Mouse | Human |
| **XCR1** | Cell Signaling<br>44665<br>Clone D2F8T<br>1/200 | Rabbit | Human |

| | | | |
|---|---|---|---|
| **CD1C** | Abcam<br>Ab156708<br>Clone OTI2F4<br>1/150 | Mouse | Human |
| **CLEC4C (CD303)** | Biotechne<br>MAB62991<br>Clone 992258<br>1/250 | Mouse | Human |
| **CSPG4 (NG2)** | Santa Cruz<br>Clone LHM 2<br>Sc-53389<br>1/500 | Mouse | Human<br>Mouse<br>Rat |
| **BCAN** | Abcam<br>Polyclonal<br>Ab111719<br>1/500 | Rabbit | Human<br>Rat |
| **GD2** | Abcam<br>Clone 14.G2a<br>Ab68456<br>1/500 | Mouse | Human |
| **Tenascin C** | Abcam<br>EPR4219<br>Ab108930<br>1/500 | Rabbit | Human<br>Mouse<br>Rat |
| **EGFR** | Cell Signaling<br>D38B1 XP®<br>4267<br>1/100 | Rabbit | Human<br>Mouse<br>Monkey |
| **EGFR VIII** | Sigma<br>Clone L8A4<br>MABC1126<br>1/200 | Mouse | Human |
| **IL-13Rα2** | Cell signaling<br>E7U7B<br>85677<br>1/200 | Rabbit | Human |

| | | | |
|---|---|---|---|
| **EPH-A2** | Cell signaling<br>D4A2 XP®<br>6997S<br>1/200 | Rabbit | Human<br>Mouse<br>Rat<br>Monkey |
| **HER2** | Cell signaling<br>Clone 29D8<br>2165<br>1/500 | Rabbit | Human<br>Mouse |
| **HES6** | Santa Cruz<br>Clone F-5<br>Sc-133196<br>1/100 | Mouse | Human<br>Mouse<br>Rat |
| **Aldh2** | ThermoFisher<br>Clone 4G6A3<br>MA5-17029<br>1/100 | Mouse | Human<br>Mouse<br>Rat<br>Non-human primate |
| **Fabp5** | Proteintech<br>Polyclonal<br>12348-1-AP<br>1/1000 | Rabbit | Human<br>Mouse<br>Rat<br>Pig |
| **PIK3R1** | Santa Cruz<br>Clone B-9<br>Sc-1637<br>1/500 | Mouse | Human<br>Mouse<br>Rat |
| **SPARC** | Proteintech<br>Polyclonal<br>15274-1-AP<br>1/1000 | Rabbit | Human<br>Mouse<br>Rat<br>Pig<br>Deer |
| **Elastin** | Abcam<br>Polyclonal<br>Ab21610<br>1/1000 | Rabbit | Human<br>Mouse |

| | | | |
|---|---|---|---|
| **Aldh1L1** | Abcam<br>Clone 3E9<br>Ab56777<br>1/500 | Mouse | Human<br>Mouse |
| **VCAM-1** | Santa Cruz<br>E-10<br>Sc-13160<br>1/500 | Mouse | Human<br>Mouse<br>Rat |
| **p-RAD50** | ThermoFisher<br>Polyclonal<br>PA5-118714<br>1/500 | Rabbit | Human<br>Mouse<br>Rat |
| **p-KAP1** | Abcam<br>EPR5248<br>Ab133440<br>1/500 | Rabbit | Human<br>Mouse |
| **p-Smad2** | Cell signaling<br>Clone 138D4<br>3108S<br>1/100 | Rabbit | Human<br>Mouse<br>Rat<br>Mink |
| **NFkB p65 (Ser536)** | Cell signaling<br>Clone (93H1)<br>3033S<br>1/100 | Rabbit | Human<br>Mouse<br>Rat<br>Monkey |
| **Calreticulin (CLRT)** | Abcam<br>Clone FMC 75<br>Ab22683<br>1/1000 | Mouse | Human |
| **CSF1R** | Abcam<br>Clone SP211<br>Ab183316<br>1/700 | Rabbit | Human<br>Rat |

| | | | |
|---|---|---|---|
| **NKG7** | Cell Signaling<br>Clone E6S2A<br>84835<br>1/100 | Rabbit | Human |
| **Granzyme A** | ThermoFisher<br>Clone 356412<br>MA5-24105<br>1/100 | Mouse | Human |
| **Fgl2** | Novus Bio<br>Clone 6D9<br>H00010875-M01<br>1/500 | Mouse | Human<br>Mouse<br>Rat |
| **SPP1** | Santa Cruz<br>Clone LFMb-14<br>sc-73631<br>1/1000 | Mouse | Human<br>Mouse<br>Rat |
| **MDK** | Santa Cruz<br>Clone A-9<br>Sc-46701<br>1/500 | Mouse | Human |
| **NCL** | Santa Cruz<br>Clone MS-3<br>Sc-8031<br>1/500 | Mouse | Human<br>Mouse<br>Rat |
| **NRC1** | R&D systems (Biotechne)<br>Clone 195314<br>MAB1850<br>1/500 | Mouse | Human |
| **PSAP** | Abcam<br>Clone 6F10<br>Ab51891<br>1/500 | Mouse | Human |

| | | | |
|---|---|---|---|
| **KLRD1 (CD94)** | Affinity Biosciences<br>Polyclonal<br>DF6773<br>1/500 | Rabbit | Human<br>Mouse<br>Rat |
| **CD56**<br>**(NCAM-1)** | Cell signaling<br>Clone E7X9M XP®<br>99746<br>1/100 | Rabbit | Human<br>Mouse<br>Rat |
| **IL-1b** | Cell signaling<br>Clone 3A6<br>12242<br>1/100 | Mouse | Human<br>Mouse |
| **Angiopoietin 2**<br>**(ANGPT2)** | ThermoFisher<br>Polyclonal<br>PA5-27297<br>1/500 | Rabbit | Human<br>Mouse<br>Rat |
| **FGFR1** | Cell signaling<br>Clone D8E4 XP®<br>9740<br>1/500 | Rabbit | Human<br>Mouse<br>Rat<br>Monkey |
| **CCL2** | Biotechne<br>Clone 23002<br>MAB2791<br>1/500 | Mouse | Human |
| **CD161**<br>**(KLRB1/NK1.1)** | Novus Bio<br>Clone PK136<br>NB100-77528<br>1/500 | Mouse | Mouse |
| **CD14** | Abcam<br>Clone 4B4F12<br>Ab182032<br>1/200 | Mouse | Human<br>Mouse |

| | | | |
|---|---|---|---|
| **CD49d** | ThermoFisher<br>Clone 9F10<br>14-0499-82<br>1/250 | Mouse | Human<br>Mouse<br>Non-human primate<br>Cynomolgus monkey<br>Rhesus monkey |
| **CD69** | GeneTex<br>Polyclonal<br>GTX37447<br>1/250 | Rabbit | Human<br>Mouse<br>Rat<br>Guinea pig |
| **TCF1/TCF7** | Cell signaling<br>Clone C63D9<br>2203S<br>1/500 | Rabbit | Human<br>Mouse |
| **L-Selectin (CD62L)** | Santa Cruz<br>Clone B-8<br>Sc-390756<br>1/1000 | Mouse | Human<br>Mouse<br>Rat |
| **IL-2R (CD25)** | GeneTex<br>Clone 1B5D12<br>GTX60792<br>1/500 | Mouse | Human<br>Mouse<br>Rat<br>Monkey |
| **NKG2A (CD159)** | Santa Cruz<br>Clone 16A11<br>Sc-53025<br>1/500 | Mouse | Mouse |
| **p-histone H3 (PHH3)** | Abcam<br>Clone mAbcam 14955<br>Ab14955<br>1/1000 | Mouse | Human<br>Mouse<br>Drosophila melanogaster |

| | | | |
|---|---|---|---|
| **Lamin B1** | Abcam<br>Clone 119D5-F1<br>Ab8982<br>1/500 | Mouse | Human<br>Mouse<br>Pig |
| **cGAS** | Santa Cruz<br>Clone D9<br>Sc-515777<br>1/100 | Mouse | Human |
| **γ-H2AX** | 1* Abcam<br>Polyclonal<br>ab11174<br>1/7000<br>2*Abcam<br>Clone 9F3<br>Ab26350<br>1/4000 | 1* Rabbit<br>2* Mouse | 1* Human<br>Mouse<br>2* Human<br>Mouse<br>Rat<br>Chinese hamster |
| **MCT4** | ThermoFisher<br>Polyclonal<br>BS-2698R<br>1/250 | Rabbit | Human<br>Mouse<br>Rat |
| **IRF-9** | Cell Signaling<br>Clone D2T8M<br>76684<br>1/250 | Rabbit | Human |
| **MBP** | Santa Cruz<br>Clone F-6<br>Sc-271524<br>1/1000 | Mouse | Human<br>Mouse<br>Rat |
| **CD209** | Santa Cruz<br>Clone DC28<br>Sc-65740<br>1/100 | Mouse | Human |

| | | | |
|---|---|---|---|
| **LAMP1** | Cell signaling<br>Clone D2D11 XP®<br>9091S<br>1/400 | Rabbit | Human<br>Monkey |
| **Rab5** | Cell signaling<br>Clone C8B1<br>3547S<br>1/200 | Rabbit | Human<br>Mouse<br>Rat<br>Monkey |
| **Rab7** | Cell signaling<br>Clone D95F2 XP®<br>9367S<br>1/200 | Rabbit | Human<br>Mouse<br>Rat<br>Monkey |
| **CD64** | ThermoFisher<br>MA5-29706<br>Clone 027<br>1/100 | Rabbit | Human<br>Mouse<br>Rat |
| **MARCO** | Abcam<br>Ab239369<br>EPR22944-64<br>1/250 | Rabbit | Human |
| **Secondary antibodies** | | | |
| **Rb_AF647** | ThermoFisher<br>Scientific<br>A32733<br>1/400 | Goat | Anti-rabbit |
| **Ms_AF555** | ThermoFisher<br>Scientific<br>A32727<br>1/200 | Goat | Anti-mouse |
| **Ms_AF647** | ThermoFisher<br>Scientific<br>A32728<br>1/400 | Goat | Anti-mouse |

| | | | |
|---|---|---|---|
| **DAPI** | ThermoFisher Scientific<br>62248<br>1/1500 | N/A | N/A |

**Supplementary Table 2: Specific CNS/glioma markers by category**

| Architectural | Cell lineage | Functional |
|---|---|---|
| ACTA2 | GFAP | Ki67 |
| CD31 | CD68 | p-STAT3 |
| | P2RY12 | HLA-DR |
| | CD11c | |
