## Supplementary Figures for "High Dimensional Proteomic Multiplex Imaging of the Central Nervous System Using the COMET™ System"

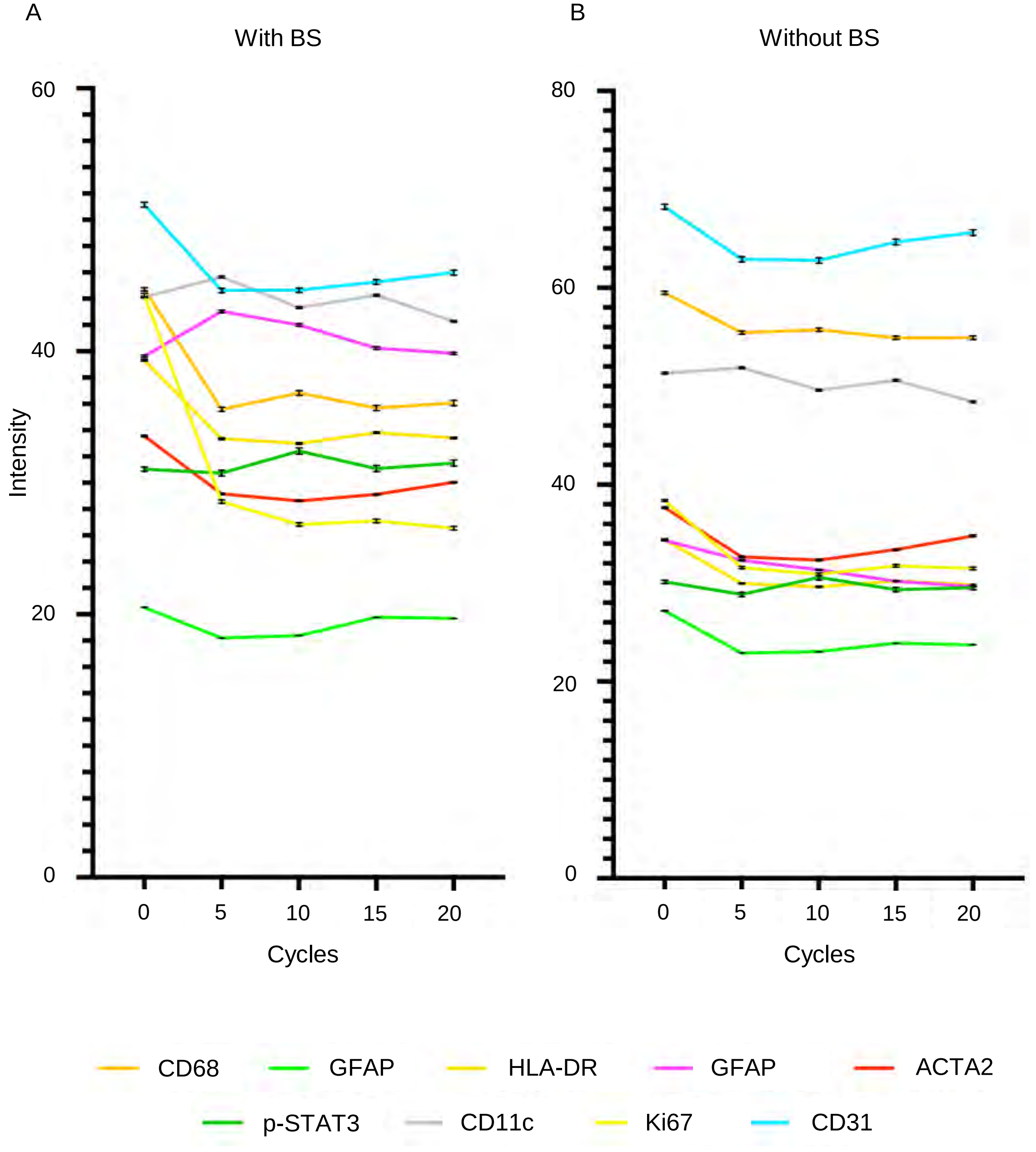
A
With BS
B
Without BS
Intensity
60
40
20
0
80
60
40
20
0
0
5
10
15
20
Cycles
0
5
10
15
20
Cycles
CD68
GFAP
HLA-DR
GFAP
ACTA2
p-STAT3
CD11c
Ki67
CD31

Suppl. Fig. 1

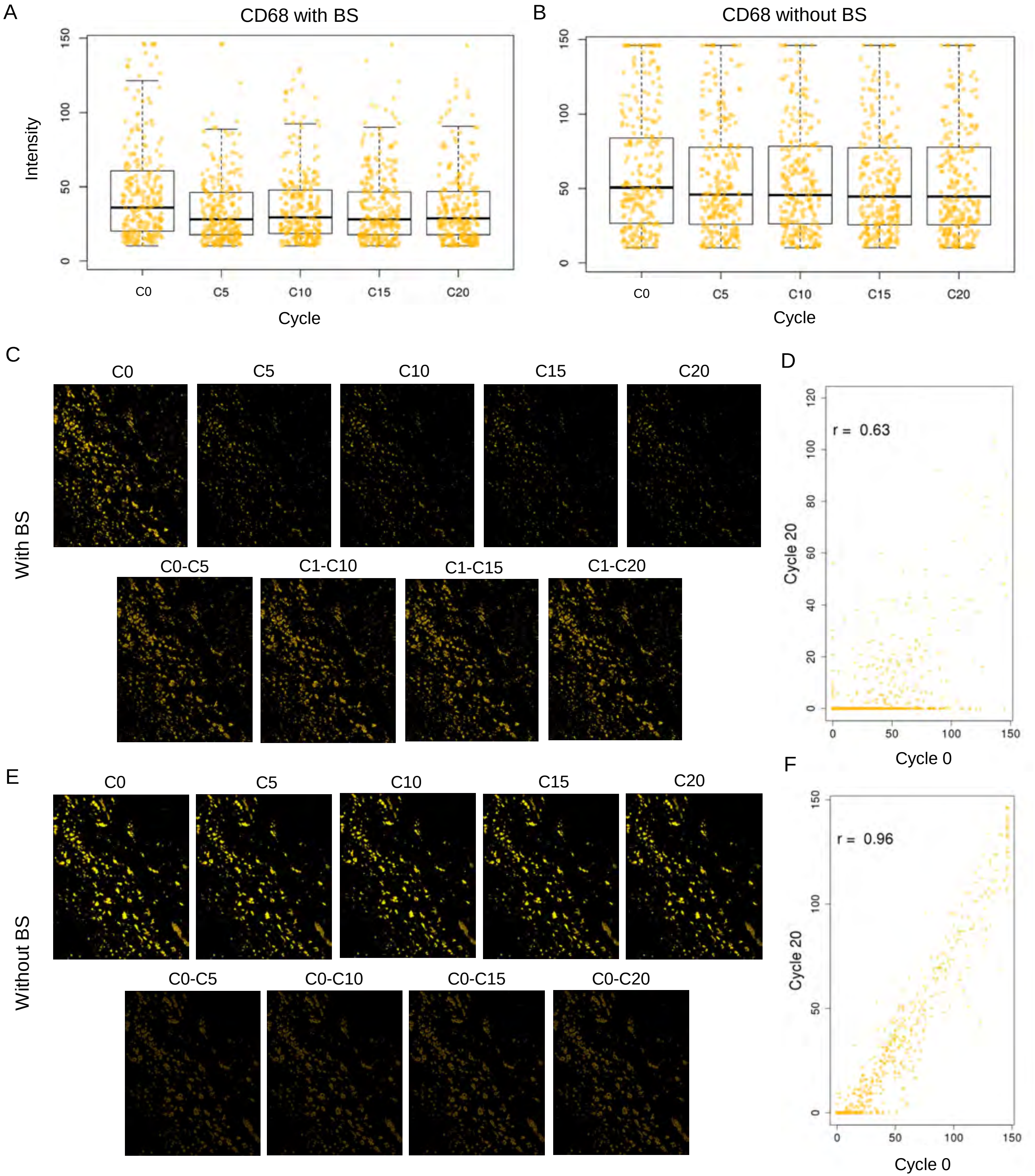

A
CD68 with BS
Intensity
0
50
100
150
C0
C5
C10
C15
C20
Cycle
B
CD68 without BS
0
50
100
150
C0
C5
C10
C15
C20
Cycle
C
With BS
C0
C5
C10
C15
C20
C0-C5
C1-C10
C1-C15
C1-C20
D
r = 0.63
Cycle 20
0
20
40
60
80
100
120
0
50
100
150
Cycle 0
E
Without BS
C0
C5
C10
C15
C20
C0-C5
C0-C10
C0-C15
C0-C20
F
r = 0.96
Cycle 20
0
50
100
150
0
50
100
150
Cycle 0


Suppl. Fig. 2

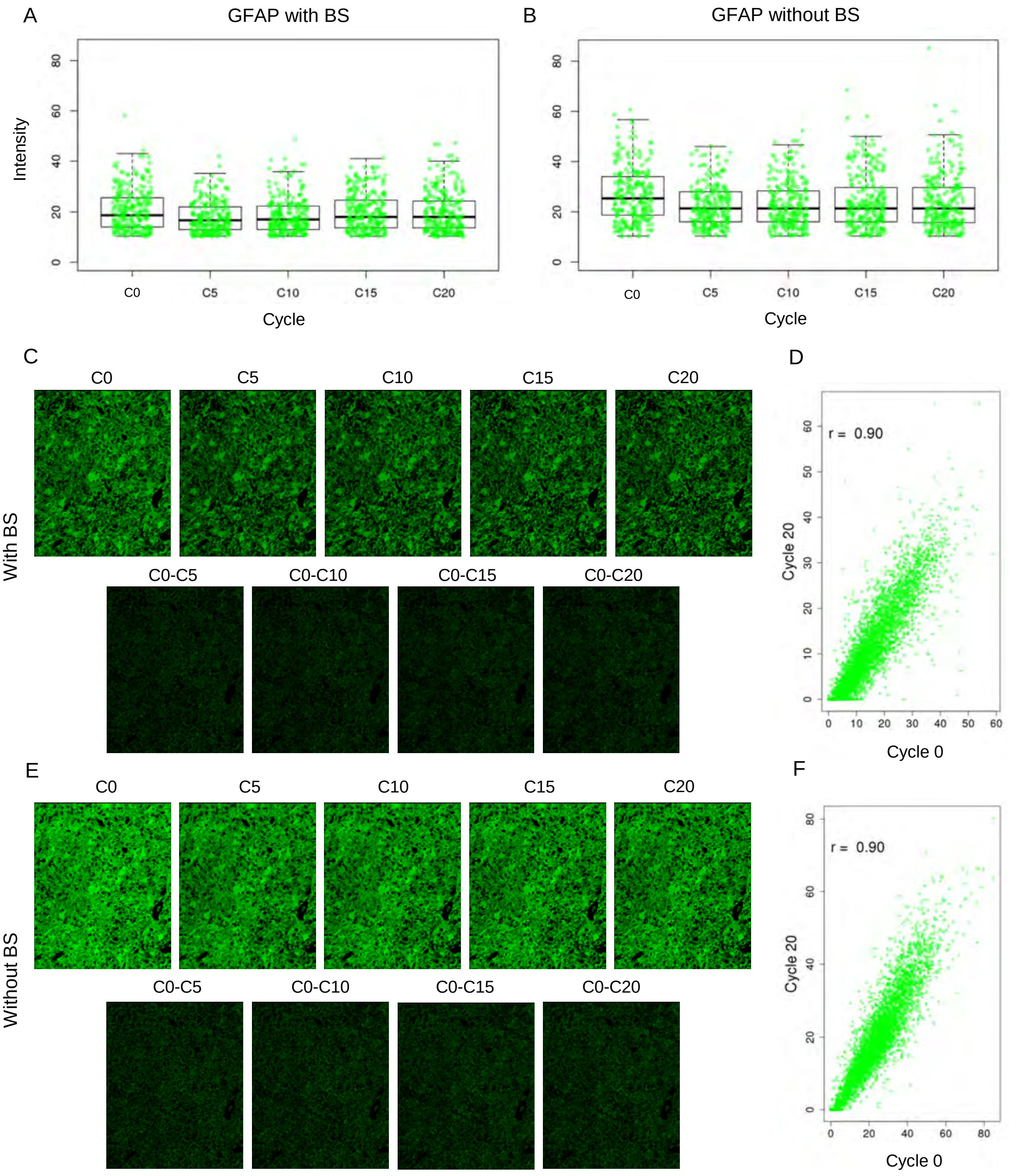

A
GFAP with BS
Intensity
0
20
40
60
80
C0
C5
C10
C15
C20
Cycle
B
GFAP without BS
0
20
40
60
80
C0
C5
C10
C15
C20
Cycle
C
C0
C5
C10
C15
C20
With BS
C0-C5
C0-C10
C0-C15
C0-C20
D
r = 0.90
Cycle 20
0
10
20
30
40
50
60
Cycle 0
E
C0
C5
C10
C15
C20
Without BS
C0-C5
C0-C10
C0-C15
C0-C20
F
r = 0.90
Cycle 20
0
20
40
60
80
Cycle 0


Suppl. Fig. 3

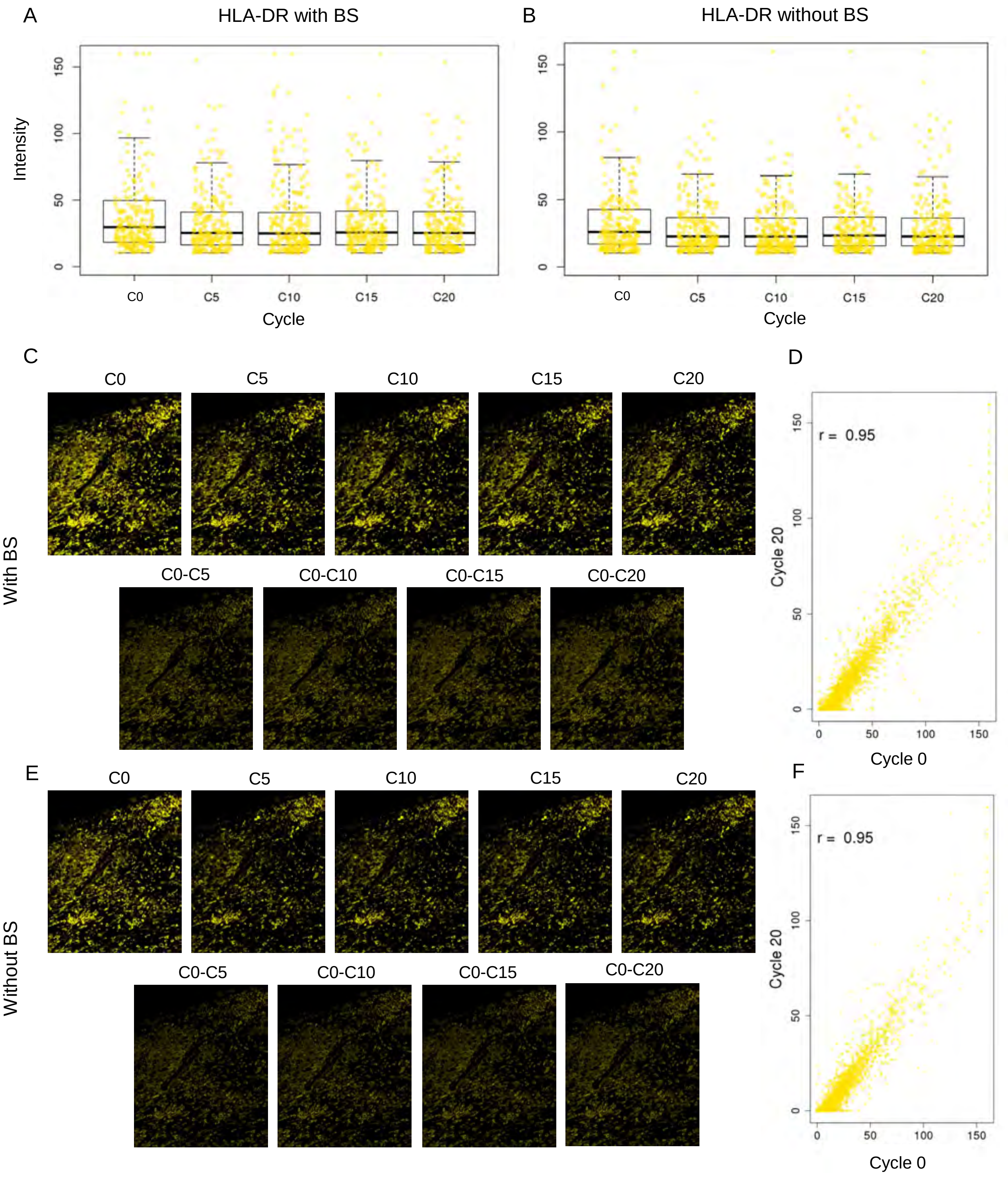

A
HLA-DR with BS
Intensity
0
50
100
150
C0
C5
C10
C15
C20
Cycle
B
HLA-DR without BS
0
50
100
150
C0
C5
C10
C15
C20
Cycle
C
C0
C5
C10
C15
C20
With BS
C0-C5
C0-C10
C0-C15
C0-C20
D
r = 0.95
Cycle 20
0
50
100
150
Cycle 0
E
C0
C5
C10
C15
C20
Without BS
C0-C5
C0-C10
C0-C15
C0-C20
F
r = 0.95
Cycle 20
0
50
100
150
Cycle 0


Suppl. Fig. 4

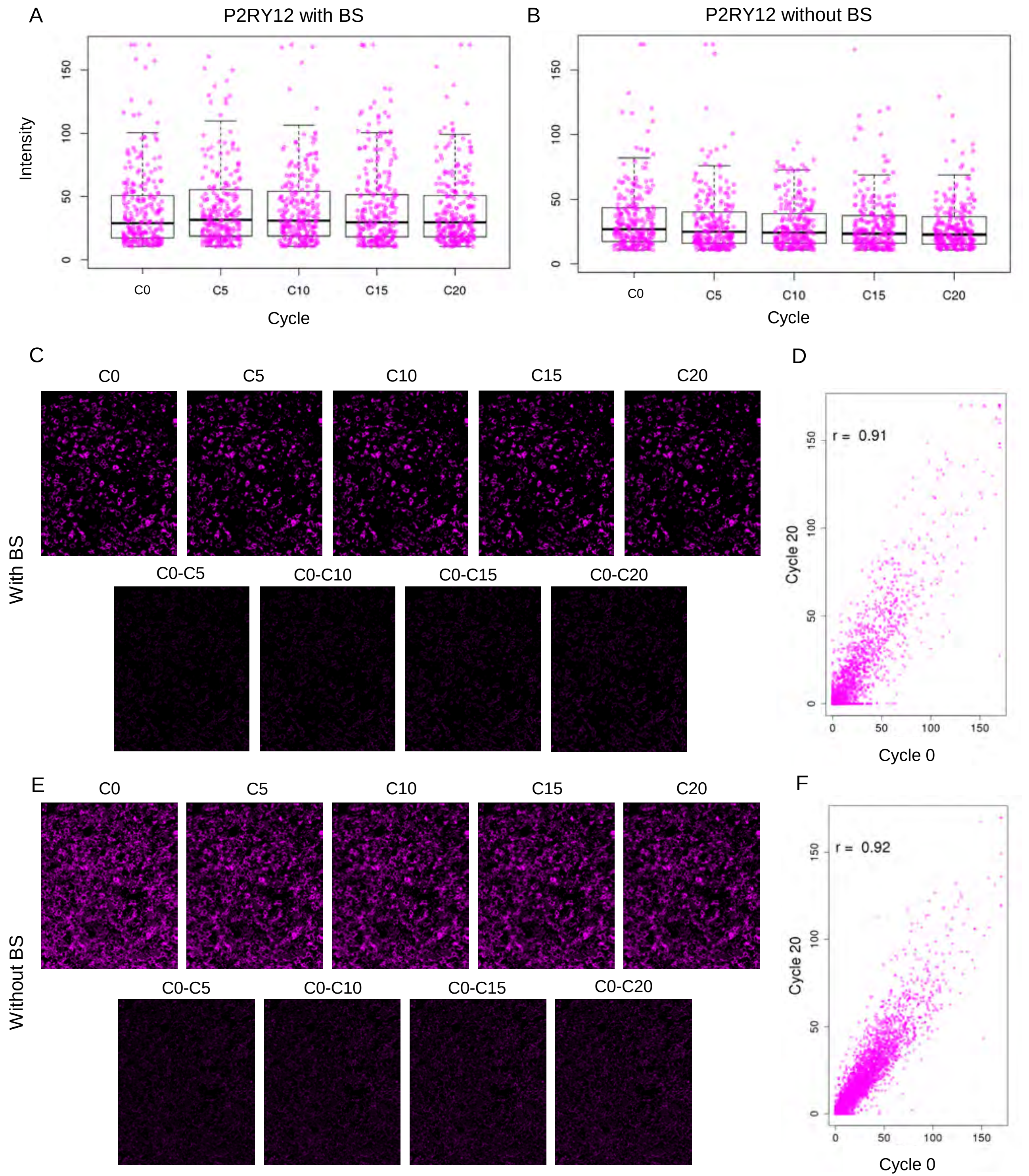
A
P2RY12 with BS
Intensity
Cycle
C0
C5
C10
C15
C20
B
P2RY12 without BS
Cycle
C
D
With BS
C0-C5
C0-C10
C0-C15
C0-C20
r = 0.91
Cycle 20
Cycle 0
E
F
Without BS
r = 0.92

Suppl. Fig. 5

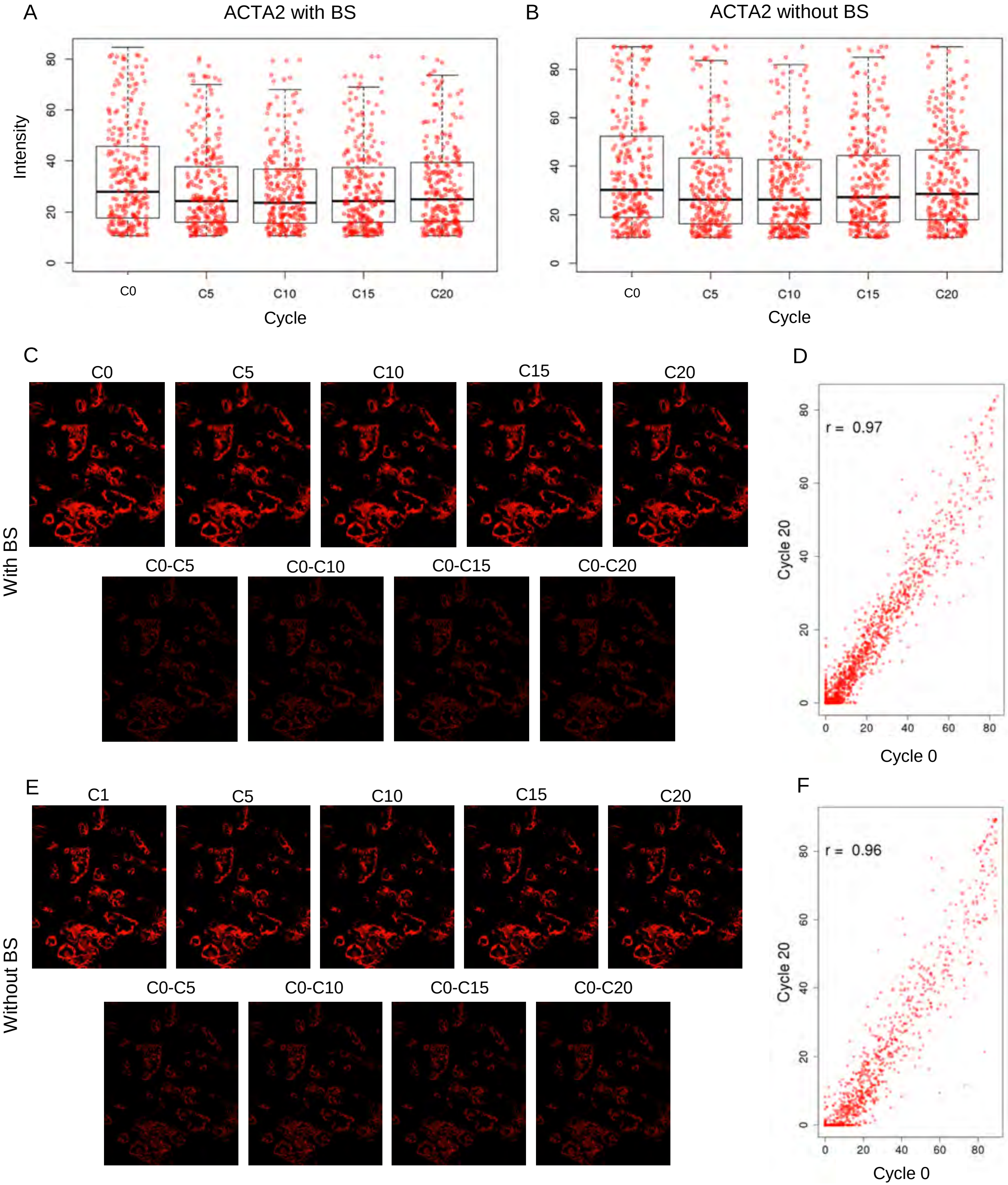
A
ACTA2 with BS
Intensity
Cycle
C0
C5
C10
C15
C20
B
ACTA2 without BS
Cycle
C
D
With BS
C0-C5
C0-C10
C0-C15
C0-C20
r = 0.97
Cycle 20
Cycle 0
E
F
C1
Without BS
r = 0.96

Suppl. Fig. 6

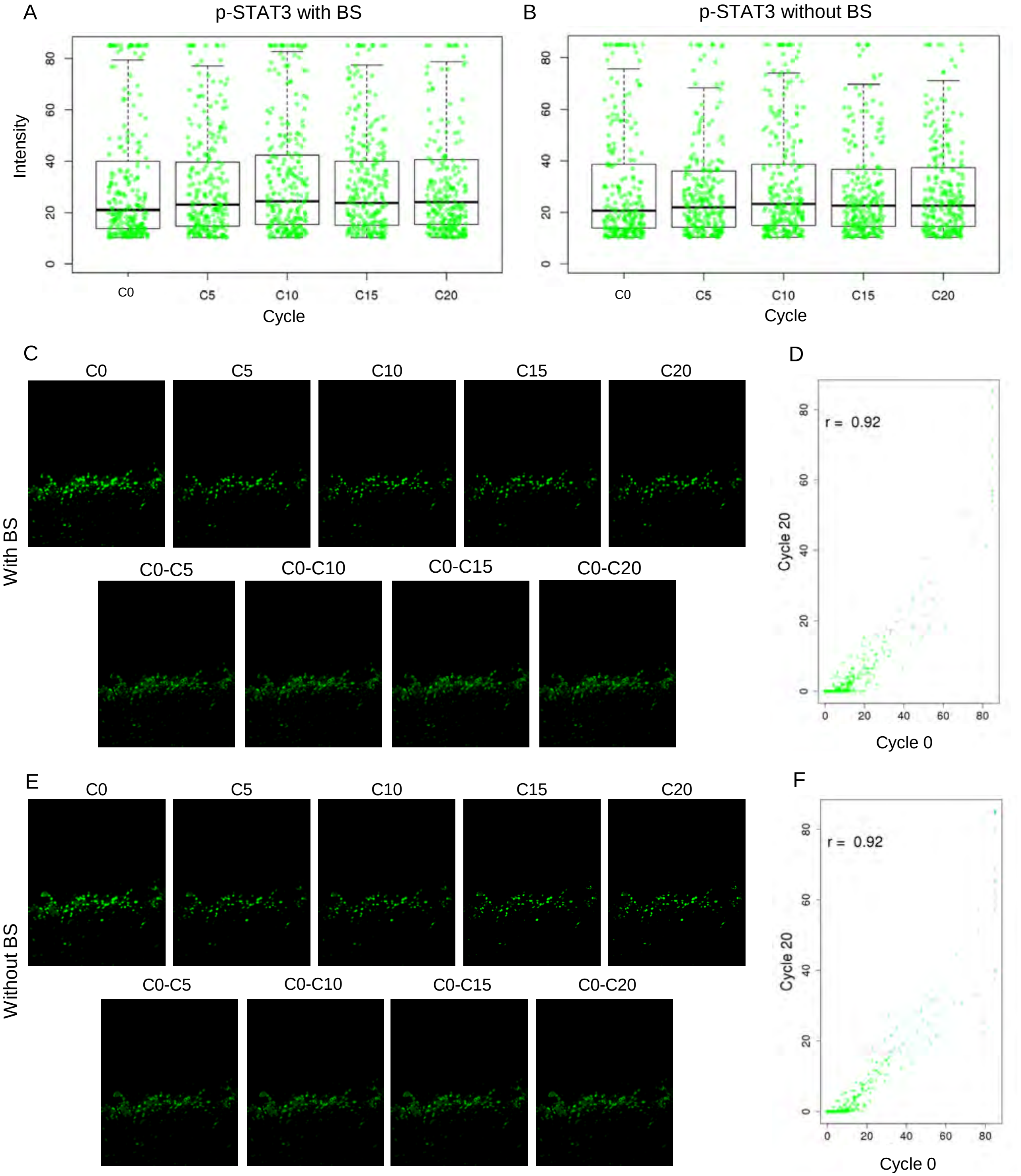
A
p-STAT3 with BS
Intensity
Cycle
C0
C5
C10
C15
C20
B
p-STAT3 without BS
C
D
r = 0.92
Cycle 20
Cycle 0
With BS
C0-C5
C0-C10
C0-C15
C0-C20
E
F
Without BS

Suppl. Fig. 7

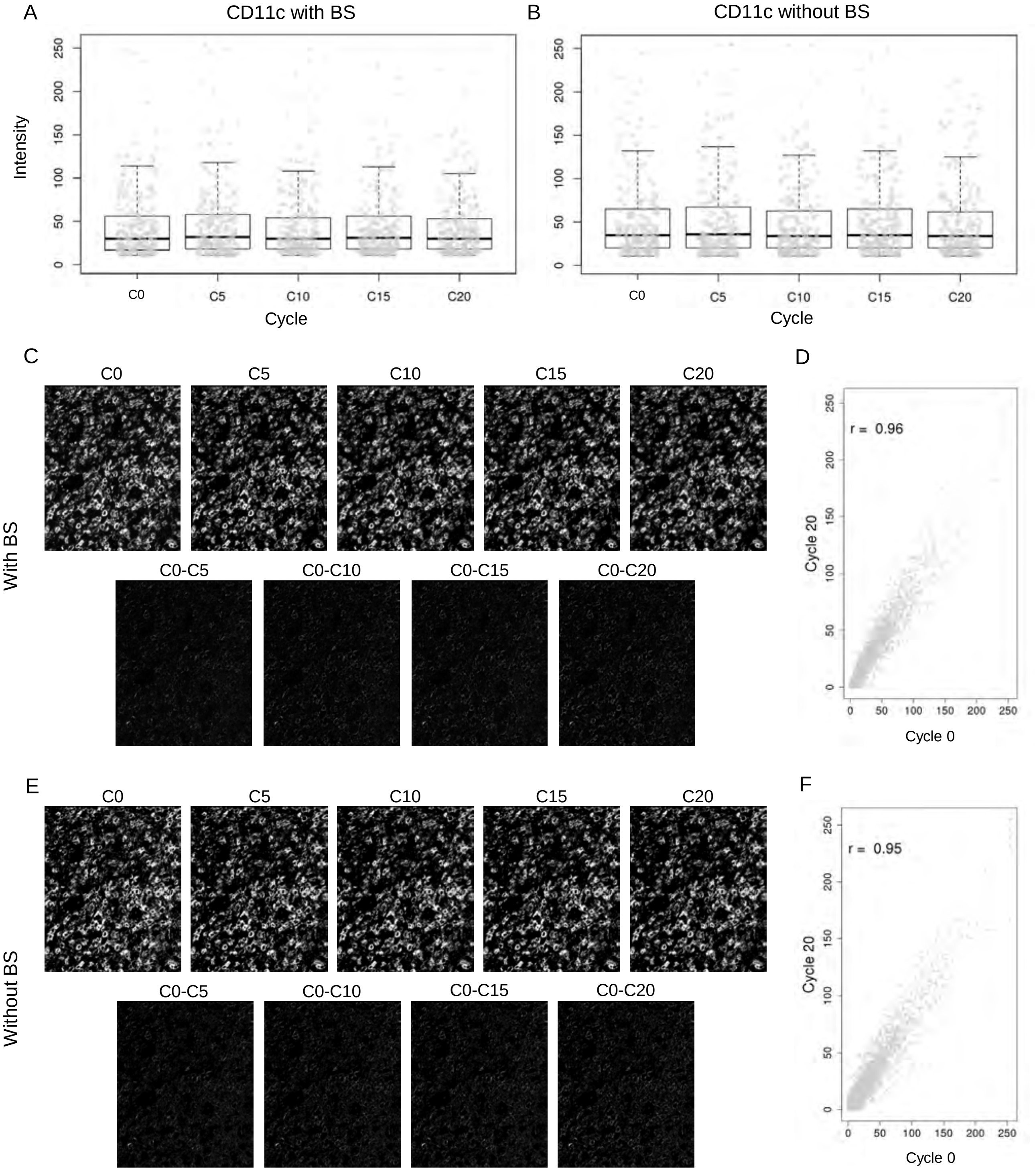
A
CD11c with BS
Intensity
Cycle
C0
C5
C10
C15
C20
B
CD11c without BS
Cycle
C
D
With BS
C0-C5
C0-C10
C0-C15
C0-C20
r = 0.96
Cycle 20
Cycle 0
E
F
Without BS
r = 0.95

Suppl. Fig. 8

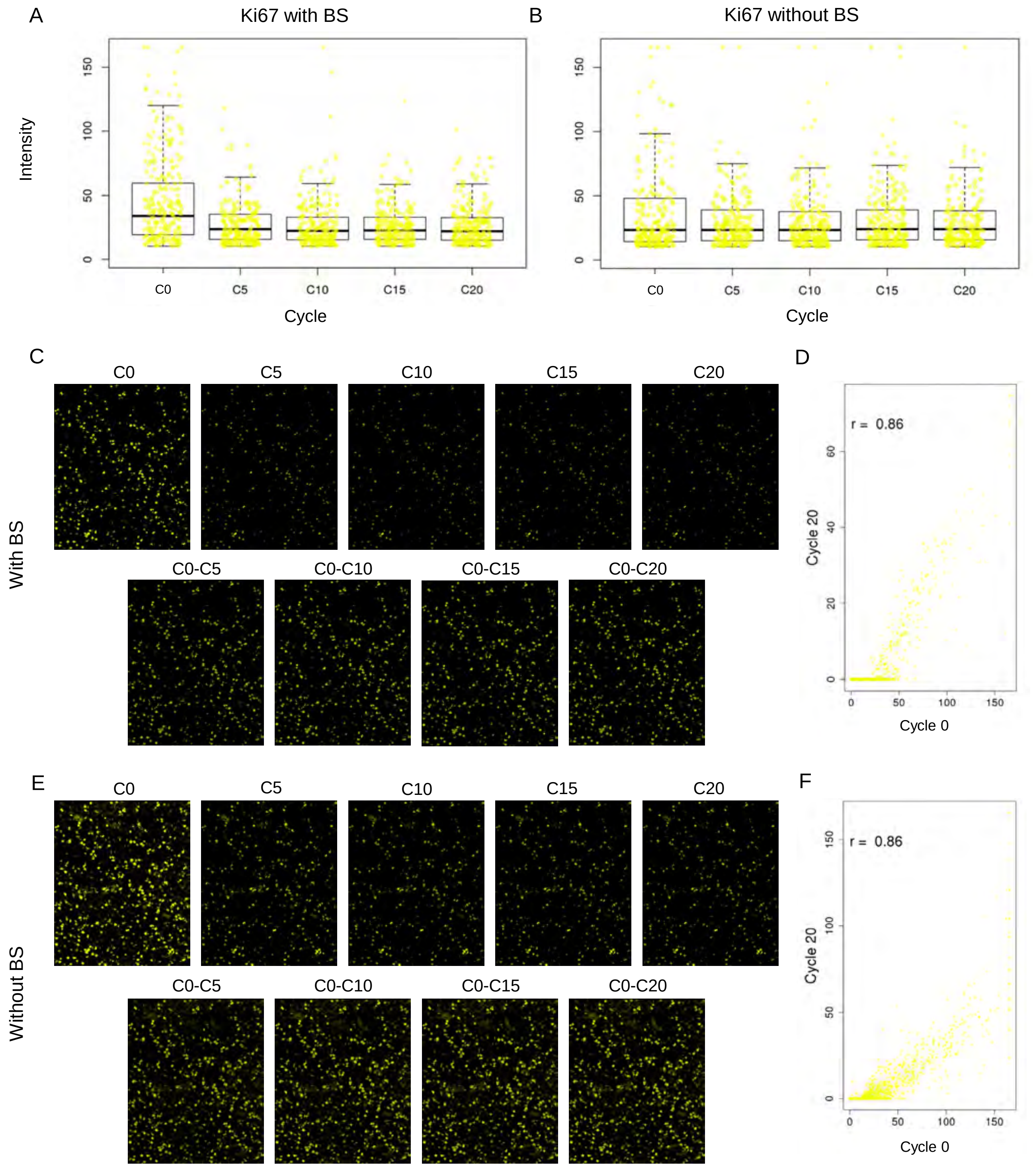

A
Ki67 with BS
Intensity
0
50
100
150
C0
C5
C10
C15
C20
Cycle
B
Ki67 without BS
C
C0
C5
C10
C15
C20
With BS
C0-C5
C0-C10
C0-C15
C0-C20
D
r = 0.86
Cycle 20
Cycle 0
E
Without BS
F


Suppl. Fig. 9

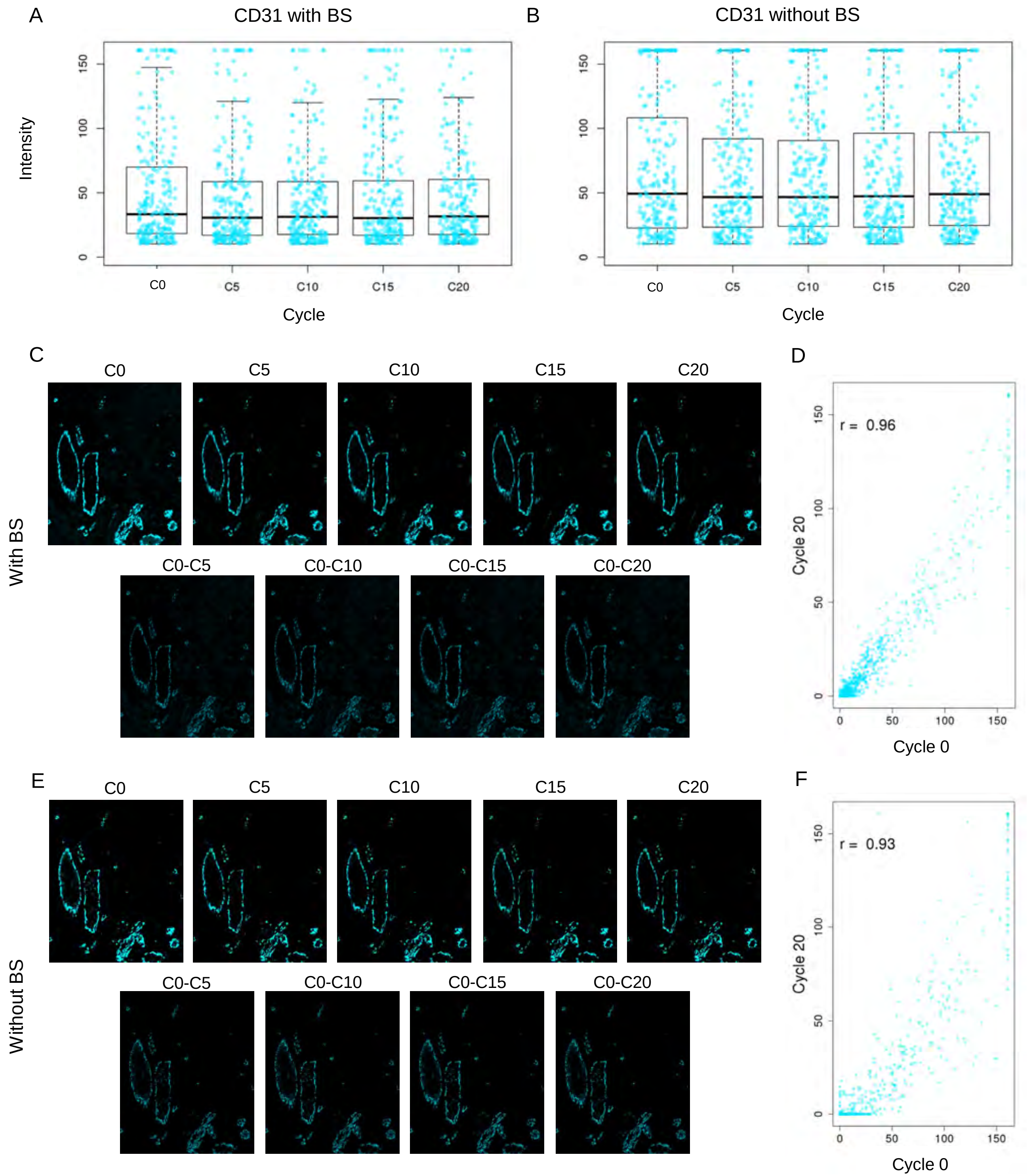
A
CD31 with BS
Intensity
0
50
100
150
C0
C5
C10
C15
C20
Cycle
B
CD31 without BS
0
50
100
150
C0
C5
C10
C15
C20
Cycle
C
C0
C5
C10
C15
C20
With BS
C0-C5
C0-C10
C0-C15
C0-C20
D
r = 0.96
Cycle 20
Cycle 0
E
C0
C5
C10
C15
C20
Without BS
C0-C5
C0-C10
C0-C15
C0-C20
F
r = 0.93
Cycle 20
Cycle 0

Suppl. Fig. 10

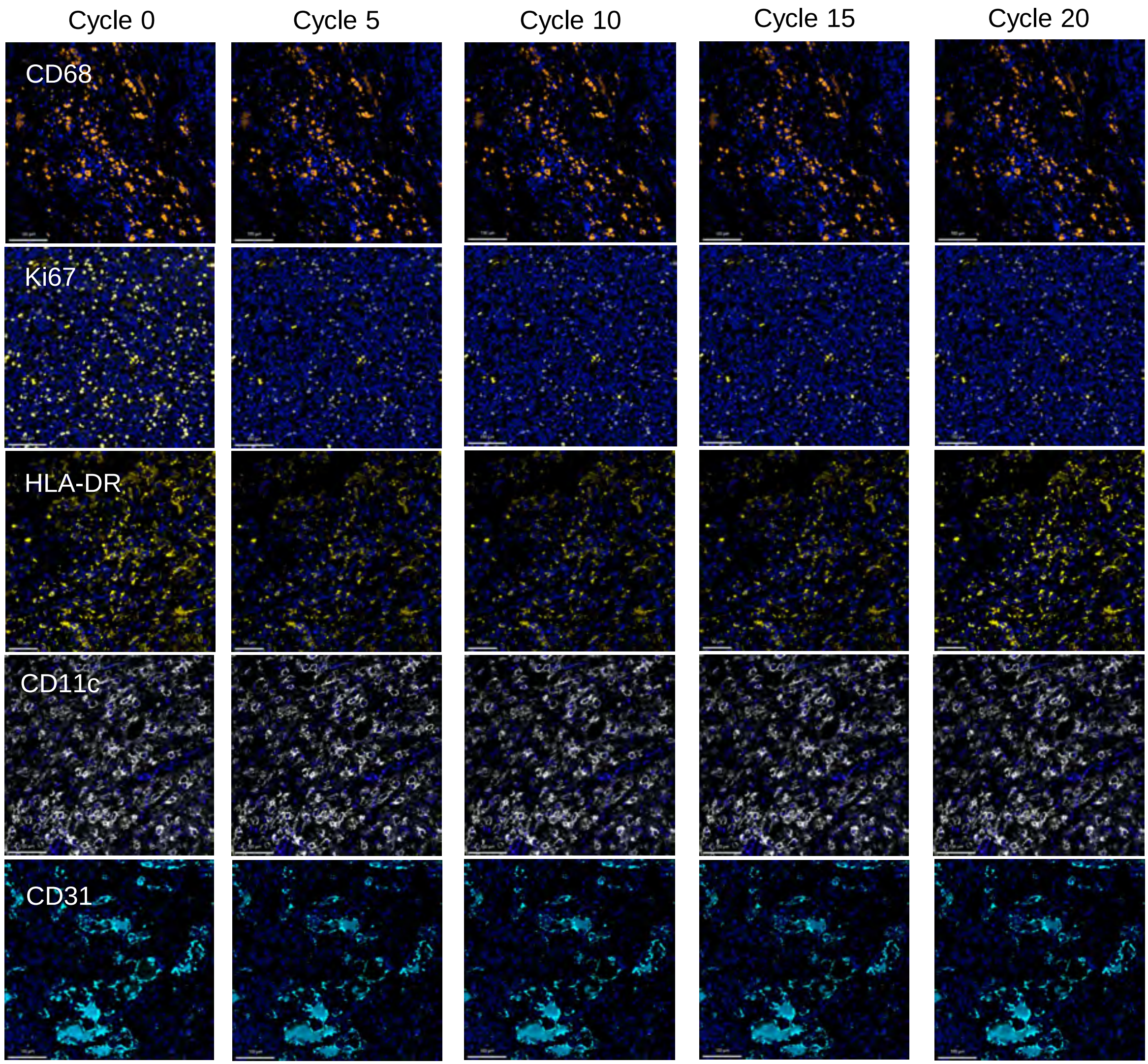

Cycle 0
Cycle 5
Cycle 10
Cycle 15
Cycle 20
CD68
Ki67
HLA-DR
CD11c
CD31


Suppl. Fig. 11

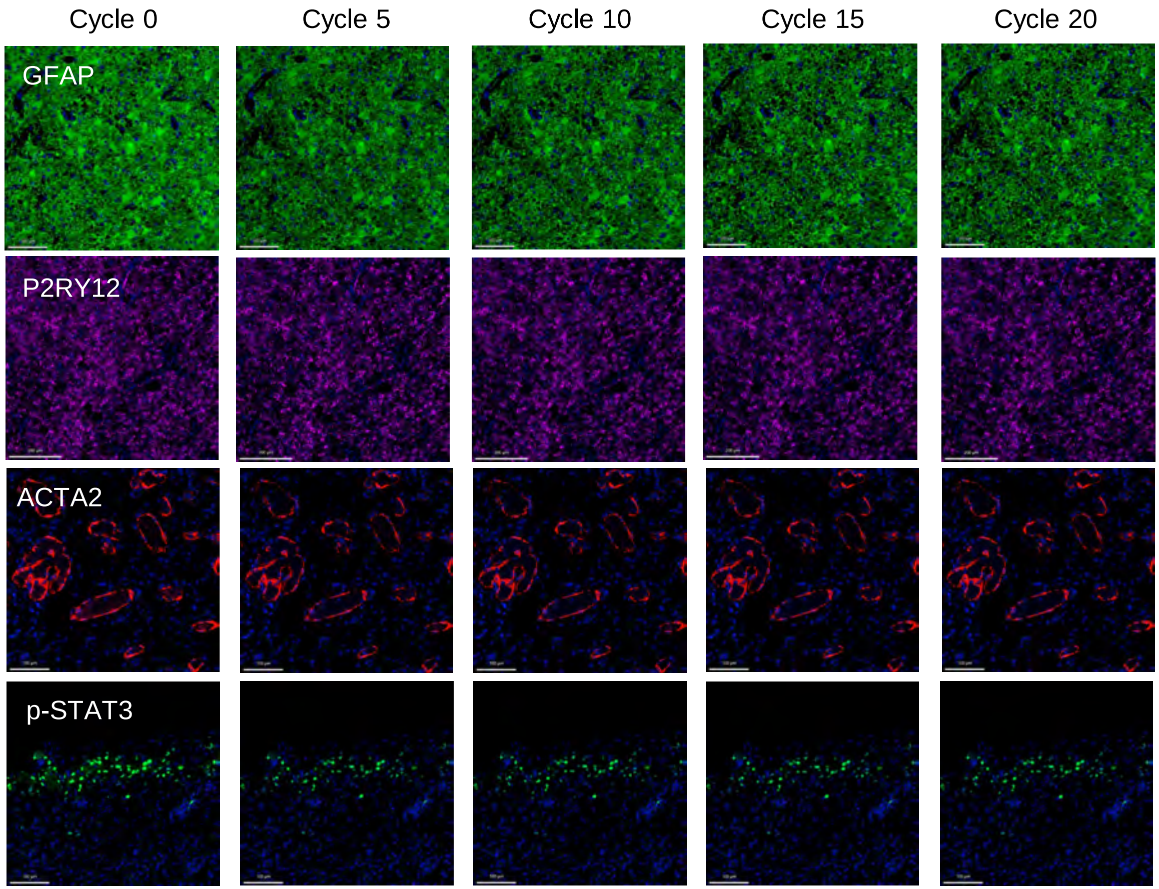
Cycle 0
Cycle 5
Cycle 10
Cycle 15
Cycle 20
GFAP
P2RY12
ACTA2
p-STAT3

Suppl. Fig. 12

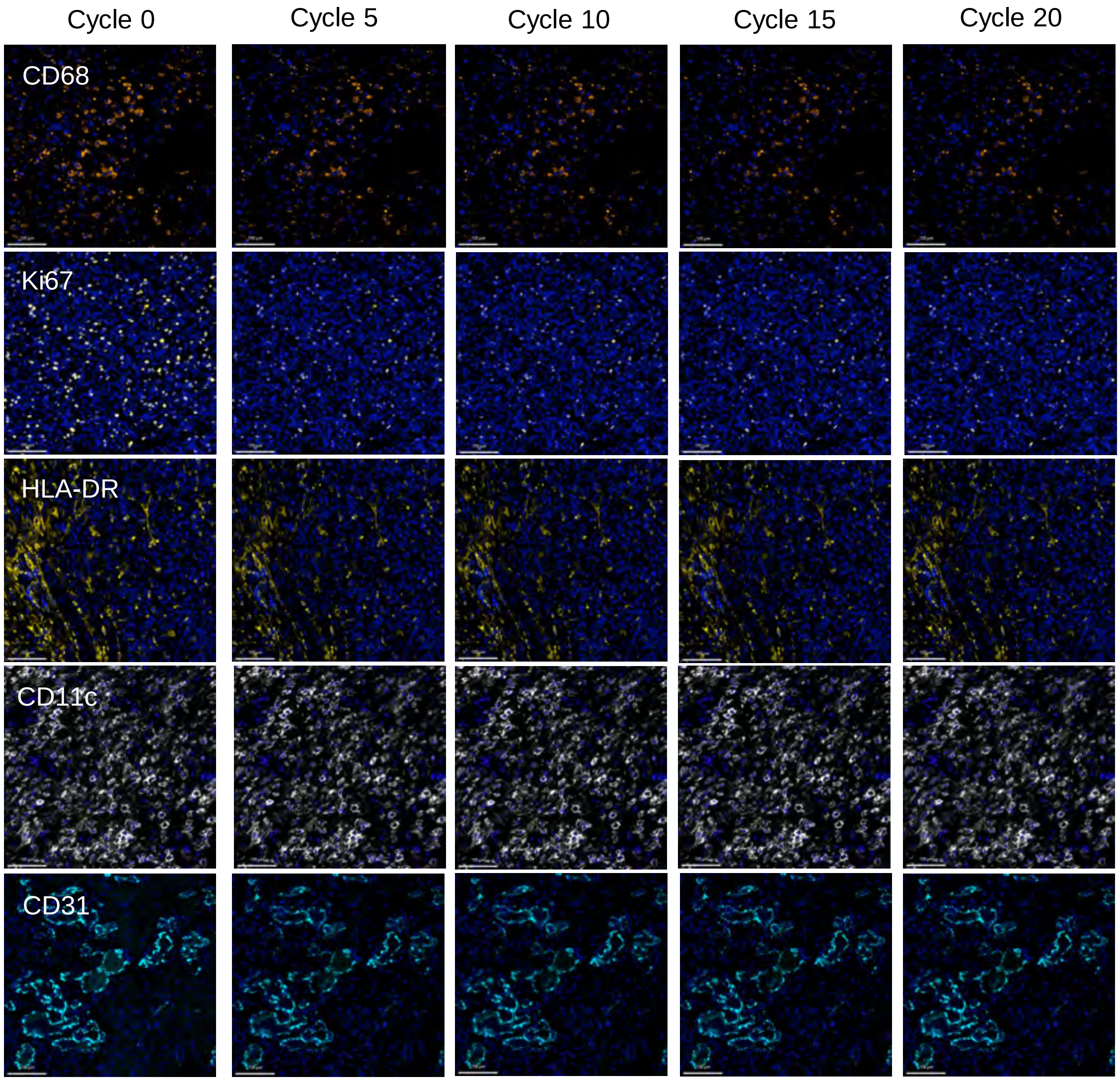
Cycle 0
Cycle 5
Cycle 10
Cycle 15
Cycle 20
CD68
Ki67
HLA-DR
CD11c
CD31

Suppl. Fig. 13

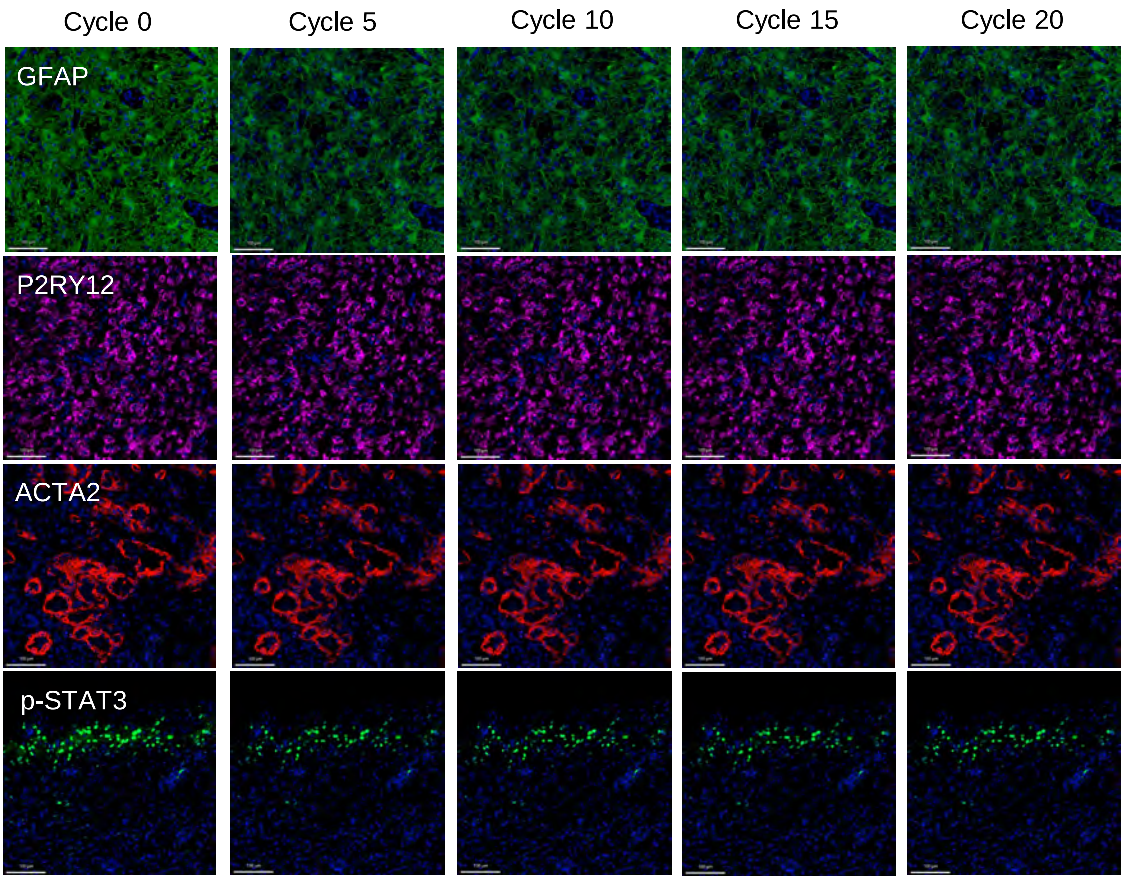
Cycle 0
Cycle 5
Cycle 10
Cycle 15
Cycle 20
GFAP
P2RY12
ACTA2
p-STAT3

Suppl. Fig. 14
